# Protein Phosphatase 2A Subunit B55 Alpha is Required for Angiotensin Type 2 Receptor Elicited Natriuresis

**DOI:** 10.1101/2025.04.07.647676

**Authors:** JJ Gildea, J Li, NL Howell, BA Kemp, MR Conaway, DL Brautigan, RM Carey, SR Keller

## Abstract

**Background:** Angiotensin type 2 receptor (AT_2_R) activation promotes natriuresis in renal proximal tubule cells (RPTCs) counteracting sodium retention stimulated by the AT_1_R. Early signaling events mediating the natriuretic response with AT_2_R activation in RPTCs are currently unknown. Our previous research suggested protein phosphatase 2A (PP2A) functions downstream of the AT_2_R. In this study, we investigated the interaction of PP2A regulatory subunit B55α with the AT_2_R and requirement for B55α in AT_2_R signaling and natriuresis.

**Methods and Results:** Rats were subjected to renal interstitial (RI) infusion of vehicle or Compound 21 (C21), a non-peptide specific AT_2_R agonist, and kidney sections were probed for interactions between PP2A subunits and the AT_2_R using a proximity ligation assay. A dramatic 6-fold increase in AT_2_R-B55α interaction in apical brush border membranes of RPTCs was observed with C21 stimulation. *In vitro* binding of purified AT_2_R and B55α supported a direct interaction between these two proteins. To determine the requirement for B55α in renal AT_2_R signaling, siRNA targeting B55α was administered to rats *in vivo* by RI infusion which resulted in a ∼70% decrease in B55α in RPTCs but not distal tubules. In rats with B55α knockdown in RPTCs, natriuresis in response to C21 was abolished. Simultaneously, C21-elicited AT_2_R redistribution to and sodium transporter Na^+^/H^+^ exchanger-3 (NHE3) retrieval from apical brush border membranes was lost, as observed with confocal immunofluorescence microscopy. Consistent with impaired AT_2_R signaling, B55α knockdown prevented c-Src phosphorylation in response to C21. B55α knockdown also led to a dramatic 4 to 6-fold increase in AT_2_R co-localization with the lysosomal marker LAMP1, and a 50% reduction in AT_2_R co-localization with EEA1 and Rab 7, markers for early and late endosomes, respectively.

**Conclusions:** PP2A B55α directly binds to the activated AT_2_R and is required for AT_2_R-elicited natriuresis, AT_2_R signaling and intracellular trafficking in RPTCs.

## INTRODUCTION

Hypertension afflicts close to half the US population^1^. Increased renal proximal tubule sodium (Na^+^) retention induced by the renal angiotensin type-1 receptor (AT_1_R) plays a major role in hypertension development^2,3^. Renal proximal tubule cells (RPTCs) also express the angiotensin type-2 receptor (AT_2_R) that inhibits Na^+^ retention and thus counteracts AT_1_R action^4,5^. Earlier studies support a critical role for AT_2_R-mediated natriuresis in blood pressure regulation as chronic renal AT_2_R activation with exogenous AT_2_R non-peptide agonist Compound 21 (C21) prevents Na^+^ retention and normalizes blood pressure in Ang II-dependent hypertension in rats^6,7^. Targeting the renal AT_2_R and its natriuretic signaling pathway may thus represent a novel option for blood pressure control. However, major gaps in our understanding of renal AT_2_R signaling hinder the development of such anti-hypertensive agents.

The AT_2_R is a seven-membrane-spanning 363 amino acid long G protein coupled receptor (GPCR)^8^. The predominant endogenous agonist for AT_2_R, at least for natriuretic responses, is angiotensin III (Ang III) that is generated after N-terminal cleavage of Ang II by aminopeptidase A^9–11^. In normal kidneys, AT_2_R activation increases Na^+^ excretion concomitant with internalization and inactivation of the sodium-hydrogen exchanger-3 (NHE3) and sodium-potassium ATPase (NKA)^6^. Translocation of AT_2_Rs from intracellular sites along microtubules to apical plasma membranes of RPTCs accompanies AT_2_R activation and natriuresis^12^. AT_2_R signaling intermediates that play roles in natriuresis include nitric oxide synthase^6^, guanylyl cyclase and cGMP^6^. The serine/threonine protein phosphatase 2A (PP2A) had previously been implicated in AT_2_R signaling in several cell types^8^, including cultured RPTCs^13^.

PP2A is a heterotrimeric enzyme composed of three subunits, A, B, and C^14^. Subunit C (36 kD) is the catalytic subunit that associates predominantly (95%) with the Aα isoform of the scaffolding A subunit (65 kD). There are four regulatory B subunit families, some with multiple spliced isoforms, B55 (α,β,γ,δ), B56 (α,β,γ,δ,ε), B72, and striatins, with B56 being the most abundant^14^. The different B subunits are structurally diverse, form distinctive heterotrimers with A and C subunits, and determine PP2A enzyme activity, substrate specificity, subcellular localization, and trafficking^15^. Recently, in experiments to uncover novel AT_2_R signaling intermediates in RPTCs that regulate natriuresis, we established that in response to renal AT_2_R activation, PP2A activity was increased^16^. Inhibition of PP2A activity with the cell permeable toxin calyculin A^17^ abolished C21-elicited natriuresis supporting the notion that PP2A activation is important for mediating natriuresis in response to AT_2_R stimulation^16^. Furthermore, we showed that PP2A subunits A, C, and B55α but not B56γ, were recruited to the apical brush border membranes of RPTCs in response to AT_2_R agonist stimulation and PP2A AB55C co-immunoprecipitated with AT_2_R^16^. Interestingly, recruitment of PP2A subunits was not observed in kidneys of pre-hypertensive spontaneously hypertensive rats (SHR) concomitant with impaired natriuresis and lack of AT_2_R redistribution to the apical brush border membrane^16^. In the current study we investigated AT_2_R and PP2A B55α interaction in RPTCs with and without AT_2_R agonist C21 stimulation *in vivo* and determined whether the two purified proteins directly bind to each other. Furthermore, we tested the requirement for the regulatory PP2A B55α subunit in AT_2_R-mediated signaling and natriuresis using RI infusion of B55α siRNA.

## METHODS

The data supporting the findings of this study are available from the corresponding author upon reasonable request.

### Animals

Experiments were conducted with 8-week-old female Wistar Kyoto (WKY) (Envigo) and 12-week-old female Sprague-Dawley (SD) (Envigo) rats. Rats were housed in a vivarium under controlled conditions (temperature 21±1°C; humidity 60±10%; 12h light/12h dark cycle) and free access to water and standard rat chow (Teklad Irradiated LM-485 Mouse/Rat Diet, cat. no. 7912). Experimental protocols were approved by the Animal Care and Use Committee of the University of Virginia and performed in accordance with the National Institutes of Health Guide for the Care and Use of Laboratory Animals.

### Experimental Procedures

#### Localization of AT_2_R Interaction with PP2A Subunits using Proximity Ligation Assay (PLA)

Female WKY rats were anesthetized with Inactin (100mg/kg) (Millipore Sigma, cat. no. T133). Jugular and carotid vessels, respectively, were catheterized for continuous infusion of 0.9% NaCl at 20 μl/min, and blood pressure measurements with a Micromed Analyzer. Rats were subjected to RI infusion of vehicle (VEH) 5% dextrose in water (D_5_W) and C21 (Cayman Chemical, cat. no. 33758), a highly selective, nonpeptide AT_2_R agonist (K_i_=0.4 nmol/l) with 25,000-fold selectivity for AT_2_Rs over AT_1_Rs, as follows. After opening the abdominal cavity open bore micro-infusion PE-10 catheters were inserted into the cortex of each kidney. Vetbond tissue adhesive (3M Animal Care Products) secured catheters and prevented interstitial pressure loss in kidneys. VEH was infused for 30 min into both kidneys at 2.5 µl/min with a syringe pump (Harvard; model 55-222). The right kidney continued to receive VEH for the remainder of the experiment while the left kidney was infused with increasing concentrations of C21 (20, 40, and 60 ng/kg/min at 2.5 µl/min, each dose for 30 min). Rats were then perfused with paraformaldehyde (PFA) as following. The left heart ventricular cavity was cannulated, and the rat perfused with 20 mL cold 4% sucrose in phosphate buffered saline (PBS) followed by 20 mL cold 4% PFA in PBS. Kidneys were dissected, cut in half, and placed in 4% PFA for 2h at room temperature. Slices were rinsed 3 x 5 min in PBS, immersed in 100 mM Tris-HCl for 30 min, and rinsed 3 x 5 min in PBS before overnight storage at 4°C in 30% sucrose in PBS. Kidney slices were embedded in Tissue Tek Optimal Cutting Tissue Compound in Cryomold vinyl specimen molds, frozen at -20°C, and stored at -80°C. Cryostat thin sections (5-8 μm) on Probe On Plus positively charged microscope slides were subjected to proximity ligation assay (PLA) using the following commercially available kits (Millipore; cat. no. DUO92102, cat. no. DUO92104 and cat. no. DUO92008-30RXN). Briefly, kidney sections were blocked in blocking buffer for 60 min at 37°C, incubated with AT_2_R antibody (Santa Cruz H-143; cat. no. sc-9040; 1:100) and simultaneously with PP2A B55α (Santa Cruz; cat. no. sc-81606; 1:100), PP2A A (Santa Cruz; cat. no. sc-13600; 1:100), or PP2A B56γ (Santa Cruz A-11; cat. no. sc-374379; 1:100) antibodies in Antibody Diluent overnight at 4°C. After washing 3 x 5 min with Wash Buffer A, kidney sections were incubated with PLA Probe for 60 min, and then in ligation buffer for 30 min at 37°C. Amplification solution was added, and slides incubated for 100 min at 37°C. After washing with 0.01X Wash Buffer B for 1 min, Fluoromount G (Southern Biotech; cat. no. 0100-01) was applied and sections covered with glass coverslips. Kidney sections were imaged using an Olympus IX81 spinning disk confocal microscope using a 60X UPlanSApo water immersion objective with a numeric aperture of 1.2. The microscope was controlled with Slidebook 5.5 software (3i, Denver CO). Five μm z-stack images at 0.25 μm intervals were captured using a Hamamatsu EMCCD camera and deconvolved using the autoquant spinning disk deconvolution module. Number of dots in designated areas (intracellular or in apical brush border membranes of renal proximal tubules) were counted and plotted. Autofluorescence identified renal proximal tubules. Specificity of AT_2_R antibodies was confirmed as previously reported^16^.

#### *In vitro* Binding Assay with Purified HA-Tagged AT_2_R and B55α

Details of this experiment are described in Supplemental Methods. Briefly, HEK293 cells were transfected with a hemagglutinin (HA)-tagged AT_2_R and cultured as described^18^. In initial experiments we observed that B55α endogenous to HEK293 cells co-immunoprecipitated with the HA-AT_2_R, and this interaction resisted washing with mild detergents. To purify the HA-AT_2_R from HEK293 cells without endogenous B55α contamination, we prepared lysates using denaturing conditions as described^19^. HA-AT_2_R was isolated from lysates with HA antibodies conjugated to agarose and incubated with 5 μg recombinant purified PP2A B55α (kindly provided by Dr. Derek J. Taylor at Case Western Reserve University) or buffer alone using a similar protocol as described^20^. Samples were analyzed by immunoblotting as described^21^.

#### Renal Interstitial Infusions of PP2A B55α siRNA and Control siRNA

Female SD rats were anesthetized using isoflurane (1.5% concentration at 3L/min). PE-10 tubing was introduced under the capsules of both kidneys and secured in place with Vetbond. Rats received RI infusion of B55α siRNA (Dharmacon Inc. Horizon Discovery Group Company, siGENOME SMARTpooL M-095376-01-0010, siGENOME Rat Ppp2r2a (117104) siRNA-SMARTpool, 10 nmol; Cat. No. D-095376-01 [CAAUAAGCCUCGUACAGUU], 02-[ACGAAUACCUCAGAAGUAA], 03-[GCUAUAUGAUGACUAGAGA], 04-[CAGUAGAGUUUAAUCAUUC]) into the left kidney while VEH (D5W) was infused into the right kidney. Another group of rats received RI infusion of scrambled siRNA (Dharmacon cat. no. K-002800-C4-01) into the left kidney while the right kidney received RI VEH infusion. siRNA (23 μg in TransIT-QR solution (Mirus Bio, Madison WI)) or VEH were infused over 30 min at 3 μL/min. PE-10 tubing was coiled and stored in a skin pouch and the skin sutured with 4-0 Prolene. Forty-eight hours later animals were anesthetized with Inactin. To facilitate breathing, PE-240 tubing was introduced into the trachea. PE-10 tubing was placed into the jugular vein to provide 1% BSA in 0.9% NaCl% at 20 µL/minute throughout the experiment. PE-50 tubing was inserted into the carotid artery to measure blood pressure every 5 min with a digital Blood Pressure Analyzer (Micromed Inc). Mean arterial pressure (MAP) was recorded for each 30-minute period. The abdominal cavity was opened to expose the right and left kidneys and ureters were cannulated using PE-10 tubing. Intrarenal PE-10 tubing left from previous RI siRNA infusions was uncoiled and used for RI infusion of VEH (D5W) into each kidney for 1h followed by infusion of increasing doses of C21 (20, 40, and 60 ng/kg/min; each for 30 min at 2.5 µl/min) into the left kidney and VEH into the right kidney. Urine was continuously collected for each 30 min time-period and urine Na^+^ excretion (UNaV) determined using a Na^+^ K^+^ Analyzer (Instrumentation Lab, model 943). At the end of the procedure, some rats were perfused with PFA and kidney sections prepared as described above for PLA. Kidney sections were used for confocal immunofluorescence microscopy for determining the degree of B55α knockdown, AT_2_R and NHE3 trafficking, and c-src signaling, and lysosomal associated membrane protein 1 (LAMP1), early endosome antigen 1 (EEA1), and small GTPase Rab7 co-localization with AT_2_R.

#### Confocal Immunofluorescence Microscopy

Kidney sections were incubated with the following single or pairs of primary antibodies in 1% milk TBST^2^ overnight at 4°C: 1) PP2A B55 (Cell Signaling; cat. no. 2290, 1:100); 2) AT_2_R (Santa Cruz H-143; cat. no. sc-9040; 1:100); 3) NHE-3 (Millipore; cat. no. MAB3136; 1:2,000); 4) p^Try416^-Src (Santa Cruz; cat. no. sc-12350; 1:100); 5) c-Src (Santa Cruz; cat. no. sc-18; 1:100); 6) AT_1_R (custom raised^22^ and kindly provided by Dr. Tahir Hussain at the University of Houston; 1:100); 7) LAMP1 (Invitrogen; cat. no. MA1-164; 1:50) and AT_2_R (Santa Cruz H-143; cat. no. sc-9040; 1:100); 8) EEA1 (Invitrogen; cat. no. 14-9114-82; 1:500) and AT_2_R (Santa Cruz H-143; cat. no. sc-9040; 1:100); 9) Rab7 (Cell Signaling; cat. no. 95746;1:100) and AT_2_R (Santa Cruz H-143; cat. no. sc-9040; 1:100). The secondary antibodies Alexa 647 conjugated donkey anti-rabbit (Invitrogen, cat. no. A31573; 1:500) or Alexa 488 conjugated donkey anti-mouse (Invitrogen; cat. nos. A31573 and A21202; 1:500), and Alexa 750 phalloidin (Invitrogen; cat. no. A30105; 1:500) in 1% milk TBST^2^ were added and incubated for 1h at room temperature. Fluoromount G was applied after washing in TBST^2^, and sections were covered with glass coverslips. Stained tubules were imaged under epifluorescence illumination as described above for PLA. RPTCs for analysis were identified based upon main cellular marker staining quality, and selected RPTCs were examined for the other antibodies as applicable. Using ImageJ software, fluorescence intensity was measured for 6 RPTCs from one rat with 4 measurements per cell averaged. To calculate immunoreactivity within RPTC apical brush border membranes, we used F-actin staining with Alexa 750 phalloidin to create a mask and determined fluorescence intensity in that region. Co-localization of the AT_2_R with LAMP1, EEA1, or Rab7 in RPTC was determined by measuring AT_2_R intracellular membrane vesicle intensity within subcellular membrane compartments marked by LAMP1, EEA1, or RAB7 as following. A proximal tubule was selected using autofluorescence in the DAPI channel (ex 350 nm/em 450 nm). Fluorescence outside the selected tubule was then cleared. Using the intensity of the subcellular vesicle marker, a mask was made using the Image/Adjust/Threshold feature and saved as an ROI file. The ROI was then opened within the AT_2_R fluorescent channel, and the Analyze Particles macro used to quantify the integrated density of AT_2_R fluorescence within each individual vesicle. The average intensity of all vesicles within one renal proximal tubule represented a single measurement. A total of 6 renal proximal tubules were analyzed and averaged.

#### Statistical Analysis

All data are presented as mean±SE. Statistical significance of quantifications for PLA and confocal immunofluorescence was determined using ordinary one-way ANOVA followed by multiple comparisons with 95% confidence using Prism Graphpad 10.2.3 (GraphPad Inc., La Jolla, CA). For Figure 4, repeated measures models using an autoregressive covariance structure were used to compare Na^+^ excretion and MAP levels. For Na^+^ excretion, the covariance matrix was chosen to reflect the double-multivariate nature of the two-kidney model, with each kidney within an animal measured under several control and C21 doses. Contrasts were used to make specific comparisons between groups and conditions. Analyses were carried out in SAS 9.4 PROC MIXED. P values <0.05 were considered statistically significant. Exact p values are given in scientific notation, but are reported as p<0.0001 when exact lower values are not provided by the statistical software. Values that were > 2 standard deviations (SD) above or below the mean were excluded from the analyses. As a result, for the 12-week-old Sprague Dawley rats used for the experiment shown in Figure 4, measures (UNaV) for control siRNA VEH, 1 rat was excluded for the second and 2 rats for the fourth experimental period, for control siRNA C21, 2 rats for C21 at 20 ng/kg/min, 1 rat at 40 ng/kg/min and 60 ng/kg/min, and for B55α siRNA C21 1 rat at 40 ng/kg/min. *In vivo* experiments were not blinded, but data analyses were blinded.

## RESULTS

### AT_2_R Interacts with PP2A B55α in Renal Proximal Tubule Cells

We previously used co-immunoprecipitation from kidney cortical homogenates to show that the AT_2_R associates with PP2A AB55αC and that the interaction increases in response to AT_2_R activation^16^. To visualize and quantify AT_2_R interactions with PP2A subunits, we performed proximity ligation assays (PLA) on kidney sections isolated from WKY rats treated *in vivo* with VEH or C21 by RI infusion. PLA is a method that detects close protein-protein interactions within a distance < 40 nm^23, 24^. Primary antibodies bound to endogenous proteins of interest are detected with oligonucleotide-conjugated secondary antibodies (PLA probes) whereby close proximity of the protein partners allows ligation of DNA strands conjugated to the secondary antibodies. The circular DNA template is amplified by PCR, detected with fluorescent DNA-binding probes, and visualized as discrete spots by confocal microscopy.

We used primary antibodies against the AT_2_R together with antibodies for PP2A subunits A, B55α, and as a control for specificity, B56γ. As shown in **Figure 1**, interactions with the AT_2_R were observed for all tested PP2A subunits. However, the localization of complexes, and changes in interactions in response to the AT_2_R agonist C21 were distinct. The number of green fluorescent dots indicating complexes of the AT_2_R with PP2A subunit B55α was relatively low with VEH treatment and predominantly localized to the intracellular space of RPTCs (**Figure 1A and B**). In response to AT_2_R activation by C21, complexes in the intracellular space of RPTCs were ∼2-fold greater (p=0.0050). However, in RPTC apical brush border membranes the number of dots increased a notable 6-fold (p=0.0125) (**Figure 1A and B**). The number of green dots representing AT_2_R interaction with PP2A A subunit were predominantly intracellular in RPTCs with VEH treatment and these were about the same after C21 (**Figure 1C and D**). However, the AT_2_R and A subunit interactions in RPTC apical brush border membranes increased ∼4.5 fold (p=0.0023) with C21 treatment (**Figure1 C and D**). There were interactions between AT_2_R and B56γ in RPTCs both intracellular and in apical brush border membranes but neither of these changed with C21 stimulation (**Figure 1E and F**). In non-RPTCs (other renal tubule segments and connective tissue and vessels surrounding tubules), signals for interactions between the AT_2_R and PP2A subunits were detected. However, the number of dots in these areas was similar for VEH and C21-treatment conditions (**Supplemental Figure 1A)**. Multiple negative controls with kidney sections undergoing PLA incubation steps with specific AT_2_R or B55α antibodies together with unspecific IgG did not yield any green dots (**Supplemental Figure 1B)**. To summarize, PLA staining of kidney sections isolated from WKY rats showed close proximity of AT_2_R with PP2A subunits. The extent of interactions with B55α and A subunits increased with *in vivo* RI C21 treatment and were most pronounced in RPTC apical brush border membranes.

**Figure 1.**
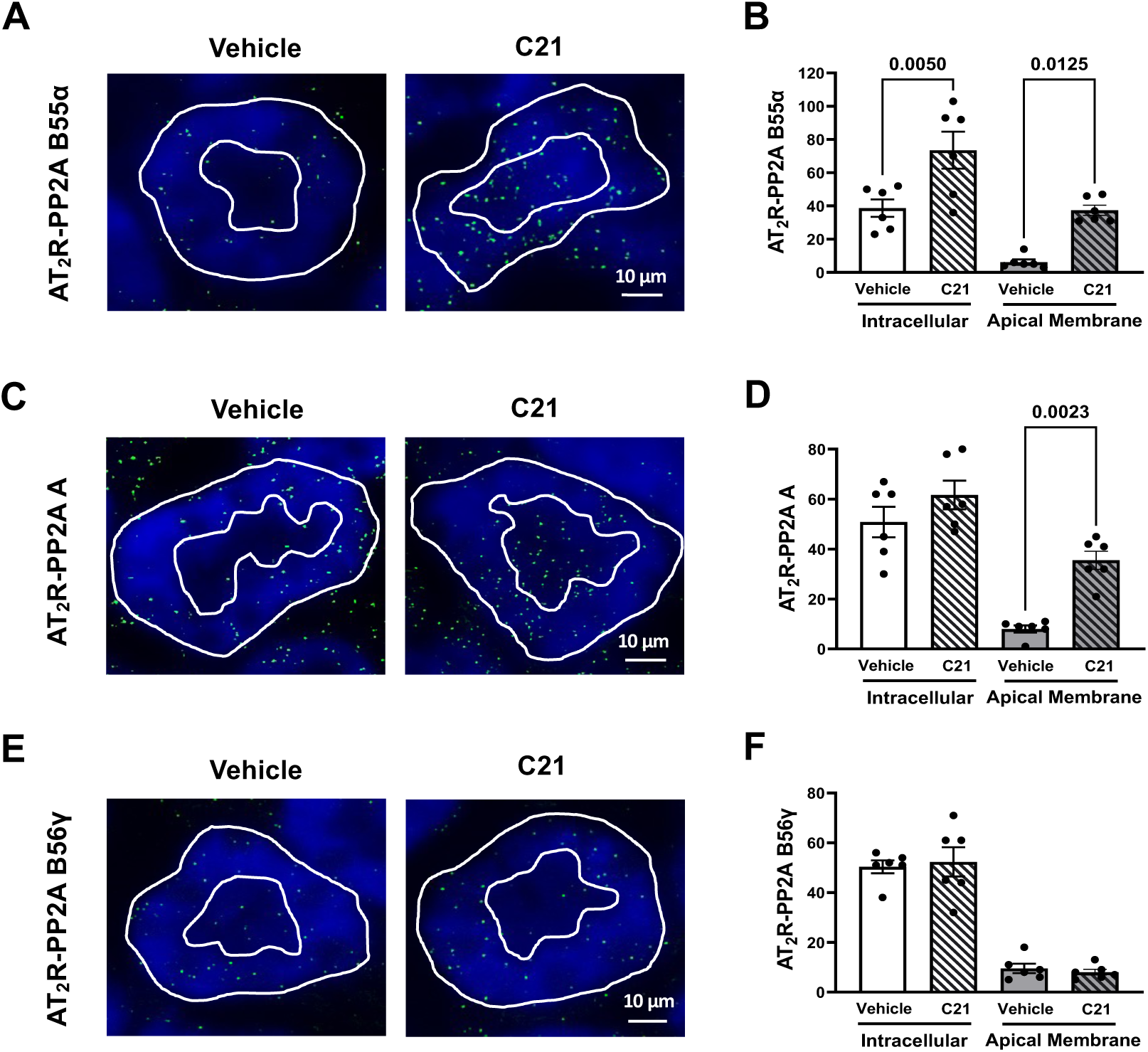
AT_2_R differentially interacts with PP2A subunits in renal proximal tubule cells (RPTC). Proximity Ligation Assays were performed on kidney sections obtained from Wistar Kyoto rats after renal interstitial infusion of vehicle or compound 21 (C21) *in vivo*. AT_2_R-specific antibodies were used in combination with antibodies against PP2A B55α **(A)**, PP2A A **(C)**, and PP2A B56γ **(E)**. Panels **A**, **C,** and **E** show representative confocal images with discrete green dots marking interactions between the AT_2_R and respective PP2A subunits. Panels **B**, **D,** and **F** show plots for the numbers of discrete green dots counted in the RPTC intracellular space (between the outer and inner white lines) and apical brush border membranes (inside the inner white line) representing interactions between the AT_2_R and PP2A B55α **(B)**, AT_2_R and PP2A A **(D)**, AT_2_R and PP2A B56γ **(F)**. Autofluorescence (blue, DAPI channel) was used to identify RPTCs. Data are shown as mean±SE (n=6) and were analyzed using one-way ANOVA.

B55α is known to mediate interaction with specific PP2A binding partners^14, 15^. To test whether AT_2_R and PP2A B55α directly interact, we purified HA-tagged AT_2_R from HEK293 cells^18^. Isolated HA-AT_2_R bound to HA-antibodies conjugated to magnetic beads was incubated with purified B55α. After incubation, beads with proteins bound to the HA-AT_2_R were pelleted, and proteins eluted from beads immunoblotted for AT_2_R and B55α. HA-AT_2_R beads that were not incubated or incubated with buffer alone served as controls (**Figure 2**, lanes 1 and 2, on left). An extract of rat kidney cortex and purified B55α served as positive controls for B55α and AT_2_R immunoblots (**Figure 2**, lanes 4 and 5). PP2A B55α was recovered in HA-AT_2_R immunoprecipitates incubated with purified B55α (**Figure 2**, lane 3) but not with HA-AT_2_R immunoprecipitates incubated with binding buffer alone (**Figure 2**, lane 2). This finding indicates direct binding of AT_2_R and B55α, consistent with their colocalization by PLA.

**Figure 2.**
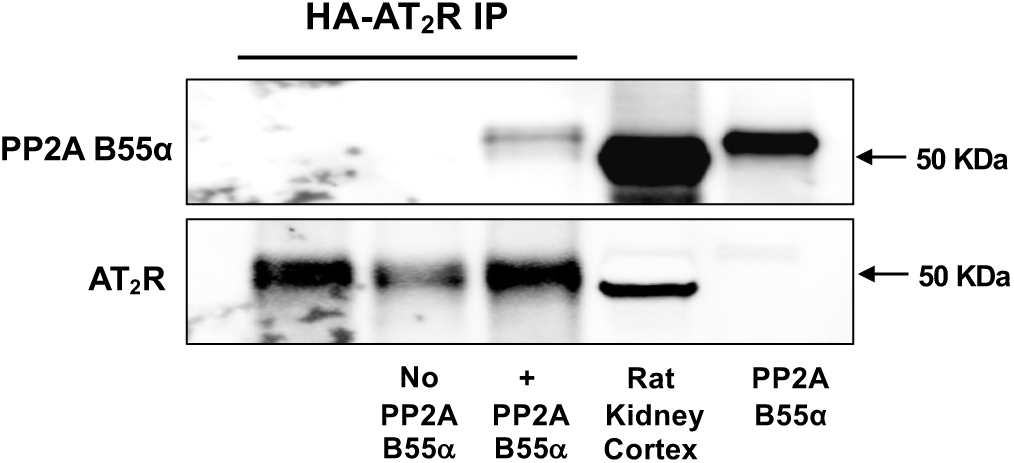
Purified AT_2_R and PP2A B55α interact in *in vitro* binding assay. HA-tagged AT_2_R was overexpressed in HEK293 cells and cell homogenates prepared under denaturing conditions to effectively disrupt protein interactions. After detergent neutralization, immunoprecipitations with HA-antibodies bound to magnetic beads pulled down HA-tagged AT_2_R (HA-AT_2_R IP). The HA-AT_2_R IP was incubated with buffer only (No PP2A B55α) or purified PP2A B55α (5 μg). Samples were separated by SDS PAGE and immunoblotted with antibodies against PP2A B55α (top panel) and AT_2_R (bottom panel). Rat kidney cortex homogenate (40 μg total protein) prepared as described^21^, and purified PP2A B55α (0.2 μg) were loaded as positive controls in right two lanes.

### Renal Interstitial Infusion of siRNA Targeting PP2A B55α Specifically Knocks Down B55α in Proximal Tubules

To test whether B55α plays a role in the natriuretic response after AT_2_R activation, we knocked down B55α by *in vivo* RI infusion of siRNA into the kidney cortex. We optimized the protocol to achieve a ∼70% (p<0.0001) reduction in B55α expression in renal proximal tubules when compared to control siRNA-infused kidneys (**Figure 3A and B**). In contrast, in renal distal tubules, where the AT_2_R is also well expressed, B55α expression was similar in B55α and control siRNA infused kidneys (**Figure 3A and B**). Concomitant with decreased total B55α expression, B55α in RPTC apical brush border membranes was decreased. In control siRNA-infused kidneys B55α in RPTC apical brush border membranes increased by ∼2-fold (p<0.0001) in response to C21 (**Figure 3C-E**). Thus, *in vivo* RI infusion of siRNA into kidney cortex is an effective approach to knock down B55α specifically in RPTCs.

**Figure 3.**
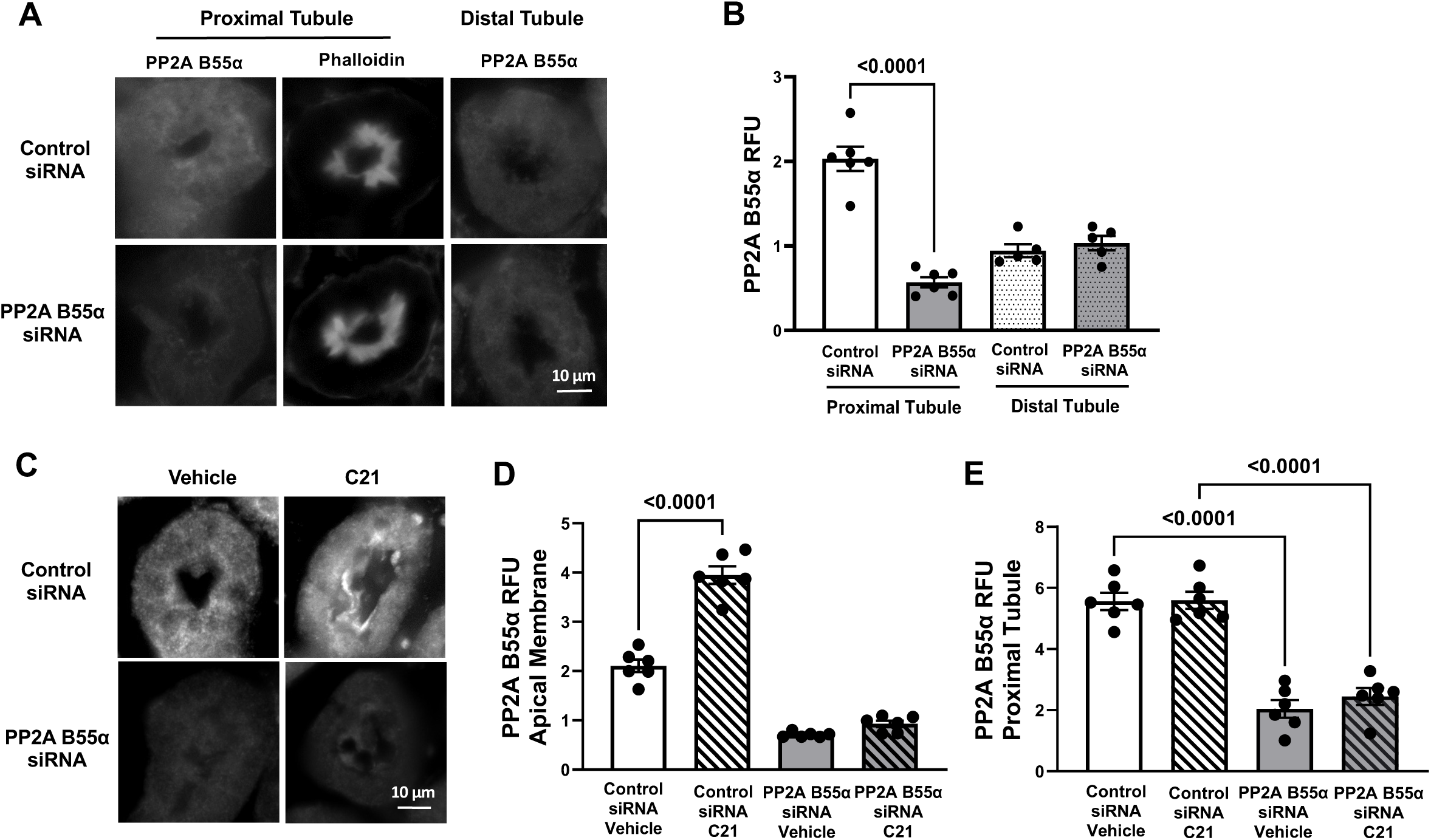
Renal interstitial infusion of siRNA targeting PP2A B55α deletes B55α in renal proximal but not distal tubule cells of Sprague-Dawley rats. **A.** Representative confocal images depict renal proximal and distal tubule staining of PP2A B55α in control siRNA and PP2A B55α siRNA treated rats 48 hours after renal interstitial (RI) infusion of siRNA (23 μg). Apical membrane staining with phalloidin 750 marks proximal tubules. **B**. Quantification of PP2A B55α fluorescence intensity (RFU) in renal proximal and distal tubule cells of control siRNA and PP2A B55α siRNA-treated kidneys. **C.** Representative confocal images of control siRNA and PP2A B55α siRNA-treated kidneys 48 hours before RI infusion of vehicle or compound 21 (C21). **D. and E.** Quantifications of renal proximal tubule cell apical membrane (D) and total (E) PP2A B55α fluorescence intensity (RFU). Data are shown as mean±SE (n=6) and were analyzed using one-way ANOVA.

### Knockdown of siRNA Targeting PP2A B55α in Renal Proximal Tubule Cells Abolishes AT_2_R-Agonist Induced Natriuresis and Impairs AT_2_R Signaling

When testing the effect of silencing B55α in RPTCs, we found no increase in urinary Na^+^ excretion (UNaV) in response to C21 in B55α siRNA-infused kidneys. In contrast, in control siRNA infused kidneys, C21 at 60 ng/kg/min stimulated UNaV ∼3.5-fold when compared to baseline or time control VEH-treated kidneys (UNaV 0.173 versus 0.049 and 0.054 μmoL/min; p<0.001) (**Figure 4A**). UNaV was similar to baseline in VEH-treated B55α- and control siRNA-treated kidneys. Mean arterial pressure (MAP) did not differ between control and B55α siRNA-RI-infused animals under all conditions (**Figure 4B**).

**Figure 4.**
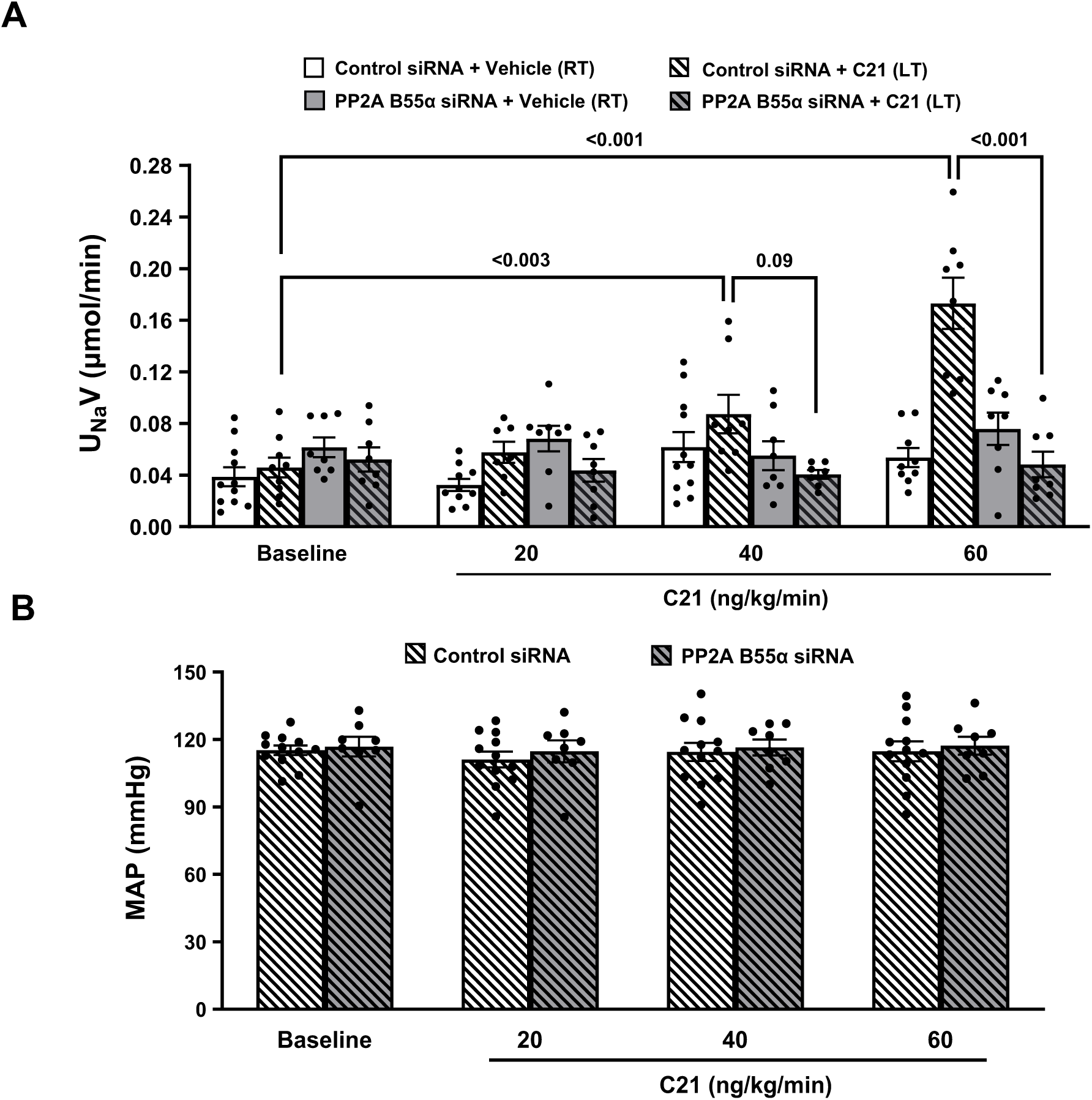
siRNA mediated knockdown of PP2A B55α in renal proximal tubule cells abolishes AT_2_R-agonist induced natriuresis. Sprague-Dawley rats were treated with control siRNA and PP2A B55α siRNA by renal interstitial (RI) infusion 48 hours prior to renal interstitial (RI) infusion of vehicle or compound 21 (C21; baseline, 20, 40, and 60 ng/kg/min, each dose for 30 minutes). Urine Na^+^ excretion (U_Na_V) (**A**) and mean arterial blood pressures (MAP) (**B**) were measured. **A.** U_Na_V measurements with the following treatments: control siRNA right kidney (RT) RI infusion of vehicle, control siRNA left kidney (LT) RI infusion of C21, PP2A B55α siRNA right kidney (RT) RI infusion of vehicle, PP2A B55α siRNA left kidney (LT) RI infusion of C21. **B.** MAP measurements in rats in which control siRNA or PP2A B55α siRNA was infused in both kidneys and vehicle into right and C21 into left kidneys. Data are shown as mean±SE. Statistical significance was determined using a repeated measures analysis with an unstructured covariance matrix in the SAS PROC MIXED program. Overall comparison for all periods between Control siRNA Vehicle (RT) and Control siRNA C21 (LT); F=30.39, P<0.001. Overall comparison for all periods between Control siRNA C21 (LT) and PP2A B55α siRNA C21 (LT); F=22.83, P<0.001.

We previously described that AT_2_R is recruited to the apical brush border membrane in response to AT_2_R stimulation and lack of recruitment is accompanied by impaired AT_2_R signaling and natriuresis^16^. When determining AT_2_R localization in siRNA-treated kidneys we found that in control siRNA/C21-treated kidneys AT_2_R increased ∼2-fold in RPTC apical brush border membranes (p<0.0001) (**Figure 5A and B**). However, B55α knockdown in RPTCs abolished C21-induced AT_2_R recruitment to apical brush border membranes of RPTCs (**Figure 5A and B**). Strikingly, with B55α knockdown AT_2_Rs localized to clearly delineated intracellular puncta in RPTCs. Total AT_2_R expression in RPTCs was similar under all conditions (**Figure 5C**).

**Figure 5.**
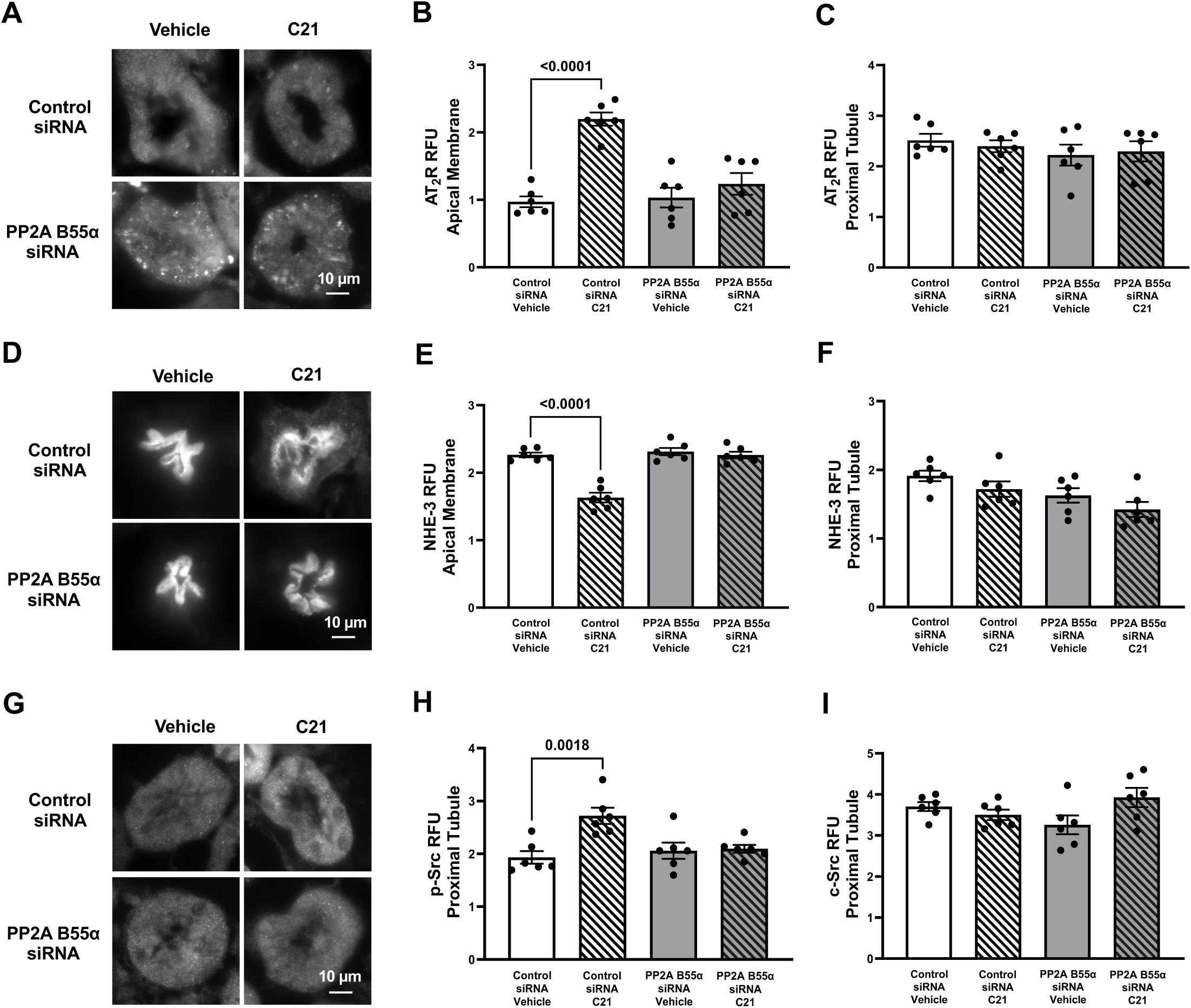
siRNA mediated knockdown of PP2A B55α in renal proximal tubule cells impairs AT_2_R signaling. Sprague-Dawley rats were treated with control siRNA and PP2A B55α siRNA by renal interstitial (RI) infusion 48 hours prior to RI infusion of vehicle or compound 21 (C21) and kidney sections analyzed by confocal microscopy analysis. Representative confocal images show proximal tubule staining for AT_2_R **(A)**, NHE-3 **(D)**, and p-Src **(G)**. Quantification of RPTC apical brush border membrane AT_2_R **(B)** and NHE-3 **(E)**, and total RPTC AT_2_R **(C)**, NHE-3 **(F)**, p-Src **(H)** and c-Src **(I)** fluorescence intensity (RFU). Data are shown as mean±SE (n=6) and were analyzed using one-way ANOVA.

NHE-3 mediates Na^+^ transport in the apical brush border microvilli of RPTCs^8^. NHE-3 retraction from this location decreases Na^+^ retention in kidney proximal tubules^8^. We previously showed that agonist-induced AT_2_R activation leads to retraction of NHE-3 from RPTC apical brush border microvilli^6^. In here, we confirm a ∼30% decrease (p<0.0001) in NHE-3 in RPTC apical brush border microvilli in control siRNA/C21-treated kidneys when compared to control siRNA/VEH treatment (**Figure 5D and E**). However, with RI infusion of B55α siRNA and C21 treatment, NHE-3 in RPTC apical brush border microvilli is the same as in VEH-treated B55α siRNA-infused kidneys (**Figure 5D and E**). Total NHE-3 expression was similar under all conditions (**Figure 5F**).

AT_2_R signaling induces c-src phosphorylation^21^. Consistent with these earlier findings, C21 induced phosphorylation of c-src on Tyr416 by 1.4-fold (p=0.0018) in control-siRNA-treated kidneys (**Figure 5G and H**). However, in B55α siRNA-treated animals, C21 failed to induce c-src phosphorylation on Tyr416. Total c-src levels were the same under all conditions (**Figure 5I**). In conclusion, knockdown of B55α abolished C21-induced natriuresis and, concomitantly, AT_2_R recruitment to and NHE-3 retrieval from RPTC apical brush border membranes, as well as intracellular signaling to c-src.

### PP2A B55α Controls AT_2_R Subcellular Distribution in Renal Proximal Tubule Cells

We noted a punctate pattern for intracellular AT_2_R staining in RPTCs that was only evident with B55α knockdown (**Figure 5A**). To determine the nature of the subcellular compartment where AT_2_Rs localized, we stained cells with antibodies against markers for select intracellular compartments; LAMP1 (lysosomal-associated membrane protein 1) for lysosomal compartments^25^, EEA1 (early endosome antigen 1) for early endosomes^26^, and the small GTPase Rab7 that controls transport to late endocytic compartments (such as late endosomes and lysosomes)^27^, and determined overlap with AT_2_R staining. AT_2_R co-localized with all 3 markers in RPTCs (**Figures 6 and 7**, **Supplemental Figures 2 and 3**), but demonstrated significant differences in the extent of co-staining in B55α siRNA and control siRNA treated kidneys. With RPTC B55α knockdown AT_2_R localization to LAMP1 positive compartments increased ∼5-fold (p<0.0001) (**Figures 6 and 7A**) while co-staining of AT_2_R with EEA1 decreased 1.7-fold (p=0.0008 and 0.0004) (**Figure 7B**) when compared to control siRNA/VEH and control siRNA/C21-treatment conditions. Interestingly, Rab7 co-localization with AT_2_R decreased 1.5-fold (p=0.0168) in response to C21 in control siRNA-infused kidneys (**Figure 7C**). With RPTC B55α knockdown, Rab7-AT_2_R colocalization was reduced by 2.9- (p<0.0001) and 2-fold (p=0.0184) reaching the same levels in vehicle and C21 treated kidneys, respectively (**Figure 7C**).

**Figure 6.**
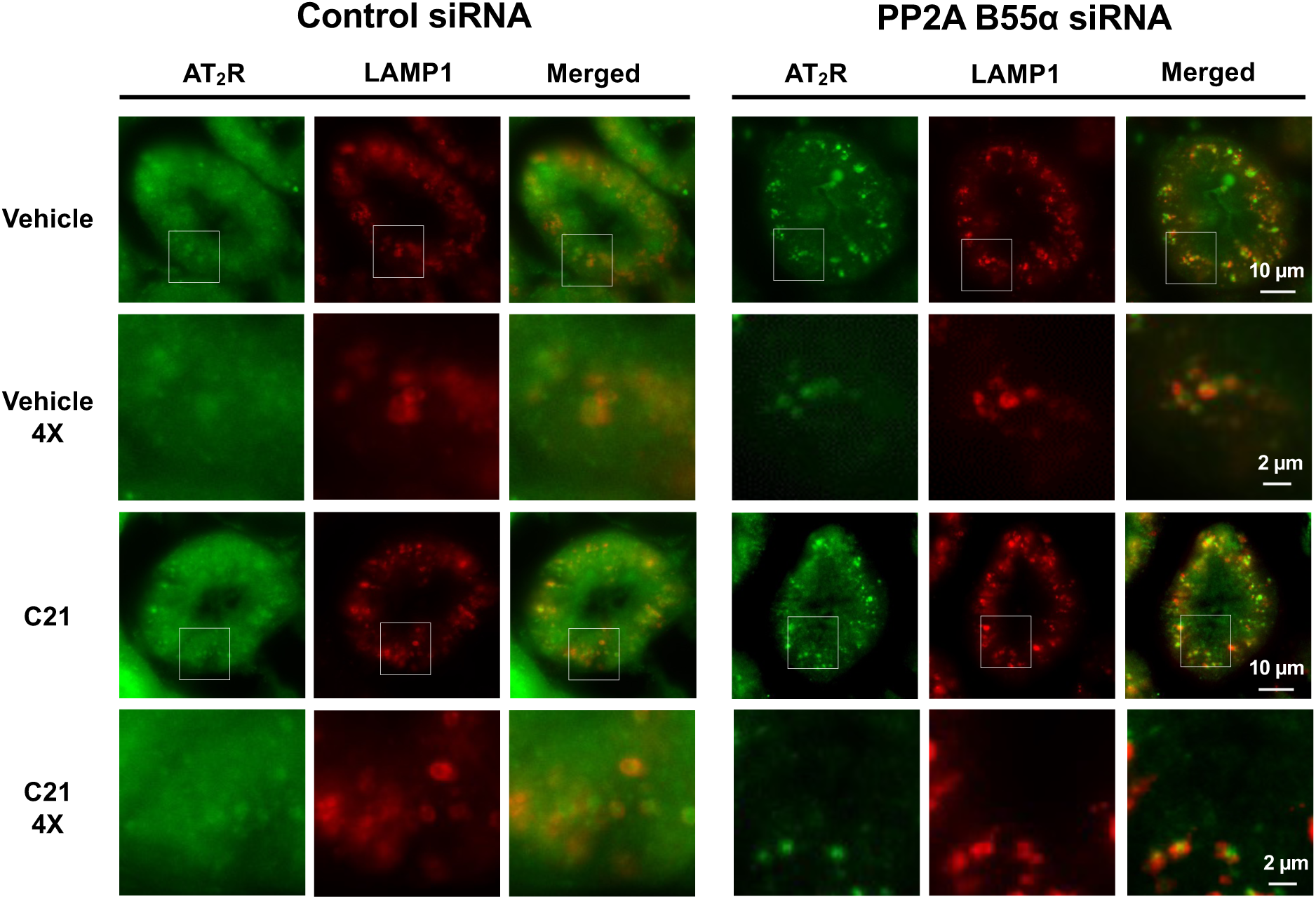
Knockdown of PP2A B55α in renal proximal tubule cells leads to accumulation of the AT_2_R in LAMP-1 positive intracellular compartments. Sprague-Dawley rats were treated with control siRNA and PP2A B55α siRNA by renal interstitial (RI) infusion 48 hours prior to RI infusion of vehicle or compound 21 (C21) and kidney sections analyzed by confocal microscopy analysis. Kidney sections were co-stained with AT_2_R and LAMP1 antibodies and representative images of renal proximal tubule cells are shown for each condition and each antibody separately and together (merged) as labeled. Squares outlined in images at 1X were magnified (4X) to better show co-localization of AT_2_R with LAMP1.

**Figure 7.**
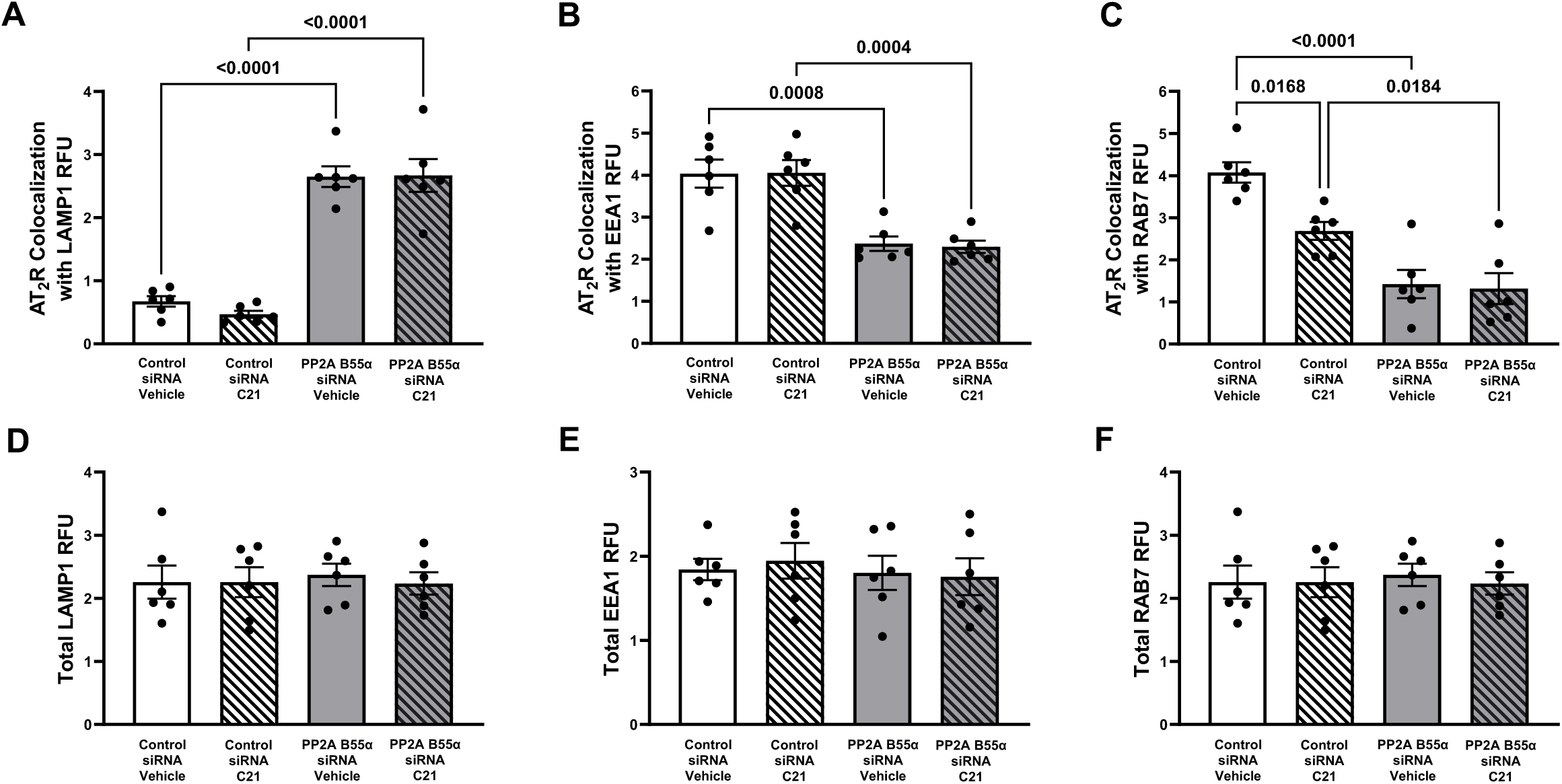
AT_2_R co-localization with LAMP1, EEA1, and Rab7 in RPTCs after treatment with control siRNA and PP2A B55α siRNA. AT_2_R co-localization with LAMP-1 **(A)**, EEA1 **(B)**, and Rab7 **(C)** marking lysosomes and early and late endosomes, respectively, was quantified in kidney sections shown in Figure 6 and Supplemental Figures 2 and 3. Total fluorescence intensity quantifications for LAMP1 **(D)**, EEA1 **(E)**, and Rab7 **(F)** in RPTCs. Data are shown as mean±SE (n=6) and were analyzed using one-way ANOVA analysis.

Total LAMP1, EEA1, and Rab7 staining was similar for all conditions (**Figure 7D-F**), as was total AT_2_R staining (**Figure 5C**). Interestingly, the AT_1_R that was previously reported to traffic through similar compartments^28^ as we describe above for the AT_2_R, maintains a normal location with B55α knockdown (**Supplemental Figure 4**). In summary, with *in vivo* RPTC B55α knockdown, AT_2_R subcellular localization shifts to a LAMP1-positive lysosomal compartment away from EEA1 and Rab7 positive early and late endosomes, respectively.

## DISCUSSION

Our results demonstrate that the AT_2_R and PP2A B55α directly interact with each other, and the interaction, visualized by PLA, increases dramatically in RPTC apical brush border membranes with AT_2_R stimulation. When B55α is knocked down *in vivo* in RPTCs, AT_2_R agonist-induced natriuresis is abolished and simultaneously AT_2_R signaling and AT_2_R redistribution to apical brush border membranes are lost. Furthermore, with B55α knockdown, AT_2_R subcellular localization shifts to a LAMP1-positive lysosomal compartment away from EEA1 and Rab7 positive early and late endosomal compartments, respectively. Our results thus support a key role for PP2A B55α in regulating both AT_2_R subcellular trafficking and signaling to natriuresis in RPTCs.

### AT_2_R Interaction with B55α and AT_2_R Signaling

The B55 and B56 family subunits mediate the binding of PP2A to selective partners, thereby determining substrate specificity and targeting. Previously identified B55α interacting proteins are involved in a wide range of cellular processes, including trafficking, cytoskeleton, and G-protein signaling^29^. Our findings indicate the AT_2_R is another protein that directly binds to B55α, and that AT_2_R stimulation strongly promotes the interaction in RPTC apical brush border membranes. This observation implies that conformational changes in the AT_2_R upon agonist binding enable binding to B55α. The 3D structure of PP2A AB55αC bound to inhibitor proteins shows that an acidic groove in B55α binds basic α-helical segments in partner proteins^30, 31^. We note that the highly conserved 43 amino acid long cytoplasmic C-terminal region of the AT_2_R has a basic α-helix^8^ similar to other B55α-interacting proteins including the GABA_B_R1 subunit of the GABA_B_ receptor, another GPCR. The B55α binding site in the GABA_B_R1 subunit was narrowed to a region in the C-terminal tail, _918_RQQLRSRRHPPT ^32^. A corresponding sequence lies in the C-terminal cytoplasmic tail of the AT_2_R, _324_RFQQKLR**S**VFRV ^8^. We speculate that B55α binding might be a property common to multiple GPCRs and the tethering may be critical to receptor signaling.

The PP2A scaffolding subunit A is also found in proximity to the AT_2_R as suggested by PLA (this study) and co-immunoprecipitation^16^. Similar to B55α, the association increases at RPTC apical brush border membranes in response to AT_2_R agonist stimulation (**Figure 1**). Since the PP2A subunit A is in a stable complex with subunit C^14, 15^, and we have previously co-immunoprecipitated the AT_2_R together with A, B55α, and C subunits^16^, we expect that AT_2_R stimulation increases AT_2_R-bound PP2A activity at RPTC apical brush border membranes. The substrate(s) for the PP2A AB55αC bound to AT_2_R is currently unknown. It is possible that the AT_2_R itself and/or other proteins recruited to the same complex serve as substrates. The aforementioned GABA_B_R1 PP2A AB55αC interaction facilitates the dephosphorylation of GABA_B_R2^32^ with which GABA_B_R1 forms a dimer. The interaction between GABA_B_R1 and PP2AB55αC was regulated by changes in intracellular calcium and the functional consequence of PP2A activity was that GABA_B_ receptor cell surface localization and signaling increased^32^.

PP2A B subunits directly affect the dephosphorylation site preference of the PP2A catalytic subunit^33^, and several possible substrates have been identified for PP2A-B55α^29, 34^ including many signaling proteins^33^. It has been reported that S/T phosphosite residues surrounded by two positively charged basic patches are more likely to undergo B55α-dependent dephosphorylation^29, 31^. This substrate recognition motif overlaps with binding partner interaction domains that B55α binds^31^. Among the five phosphorylation sites that were identified by phosphoproteomics (PhosphoSitePlus) in the AT_2_R C-terminal cytoplasmic domain^35^, the Ser331 phosphosite, detected in Jurkat cells after calyculin A and pervanadate treatment^36^, lies within the aforementioned AT_2_R basic α-helix, the potential B55α interaction domain. Phosphorylation of the AT_2_R helix might disrupt binding to B55α and thereby act as a switch to modulate complex formation. The kinase and phosphatase for Ser331 are unknown but the site is surrounded by basic residues making it an excellent candidate substrate for PP2A-B55α. Earlier studies support a role for AT_2_R dephosphorylation in AT_2_R signaling. Increased AT_2_R phosphorylation in SHR kidneys was observed with impaired C21-induced natriuresis, and natriuresis was restored with decreased AT_2_R phosphorylation^37^. We previously showed that in RPTCs of SHR, AT_2_R interaction with PP2A AB55αC is impaired with simultaneous loss of AT_2_R signaling and natriuresis^16^.

### Roles of B55α in AT_2_R Subcellular Trafficking

Little is currently known about AT_2_R subcellular trafficking, but it seems different from other GPCRs. GPCRs, including the AT_1_R, undergo rapid internalization upon ligand binding to terminate receptor signaling^28^. Earlier studies suggested that the AT_2_R, despite being a GPCR, is not internalized^38^. This was established through studies in HEK293 cells in which the AT_2_R and AT_1_R were overexpressed and cell surface expression, internalization, and recycling compared between the two receptors^38^. Our earlier *in vivo* studies in RPTCs are not consistent with static AT_2_R localization to the cell surface as a large portion of AT_2_R staining is intracellular, and AT_2_R increases in apical brush border membranes in response to AT_2_R stimulation^6, 7, 11, 16, 21^. With our co-localization studies in RPTCs of control siRNA-VEH treated RPTCs, we identify intracellular compartments to which the AT_2_R localizes under basal and agonist-stimulated conditions as EEA1- (early endosomes), Rab7- (late endosomes), and LAMP1-positive compartments (lysosomal). The AT_1_R traffics through similar compartments. It is internalized through clathrin-coated pits and caveolin-dependent pathways and moves to early endosomes from which it is recycled back to the cell surface^28^. Sustained binding of the AT_1_R to β-arrestin induces trafficking to late endosomes and lysosomes for degradation^28^. Rab7 overexpression also increases AT_1_R lysosomal degradation^39^. However, we show in here that AT_1_R subcellular localization is not affected by B55α knockdown supporting differential regulation of AT_1_R and AT_2_R subcellular trafficking. Also, the control of their cell surface localization in response to stimuli clearly differs. While agonist binding to the AT_1_R reduces cell surface localization^28^, agonist binding to the AT_2_R increases cell surface localization^6^. The divergent response may be achieved through differential phosphorylation/dephosphorylation events. AT_1_R desensitization is mediated by G protein-coupled receptor kinases (GRKs) that phosphorylate the receptor to allow recruitment of β-arrestins and promote receptor internalization through clathrin-coated pits^28^. Based on our observation that the AT_2_R does not increase in RPTC apical brush border membranes in response to AT_2_R stimulation in the absence of B55α, we suggest that dephosphorylation events triggered by AT_2_R agonist binding and B55α-AT_2_R interaction promote AT_2_R cell surface localization by decreasing AT_2_R endocytosis and/or promoting AT_2_R recycling to the cell surface (**Figure 8**). In support of dephosphorylation keeping the AT_2_R at the cell surface, we recently observed that AT_2_Rs overexpressed in HEK293 cells tightly bind the B55α subunit (unpublished) thereby possibly promoting AT_2_R and/or associated protein dephosphorylation and AT_2_R cell surface localization as previously observed^38^. Interestingly, the clathrin adaptor proteins AP1 and AP2 are substrates for PP2A-B55α and the phosphorylation status of AP2 alters endocytosis^40^.

**Figure 8.**
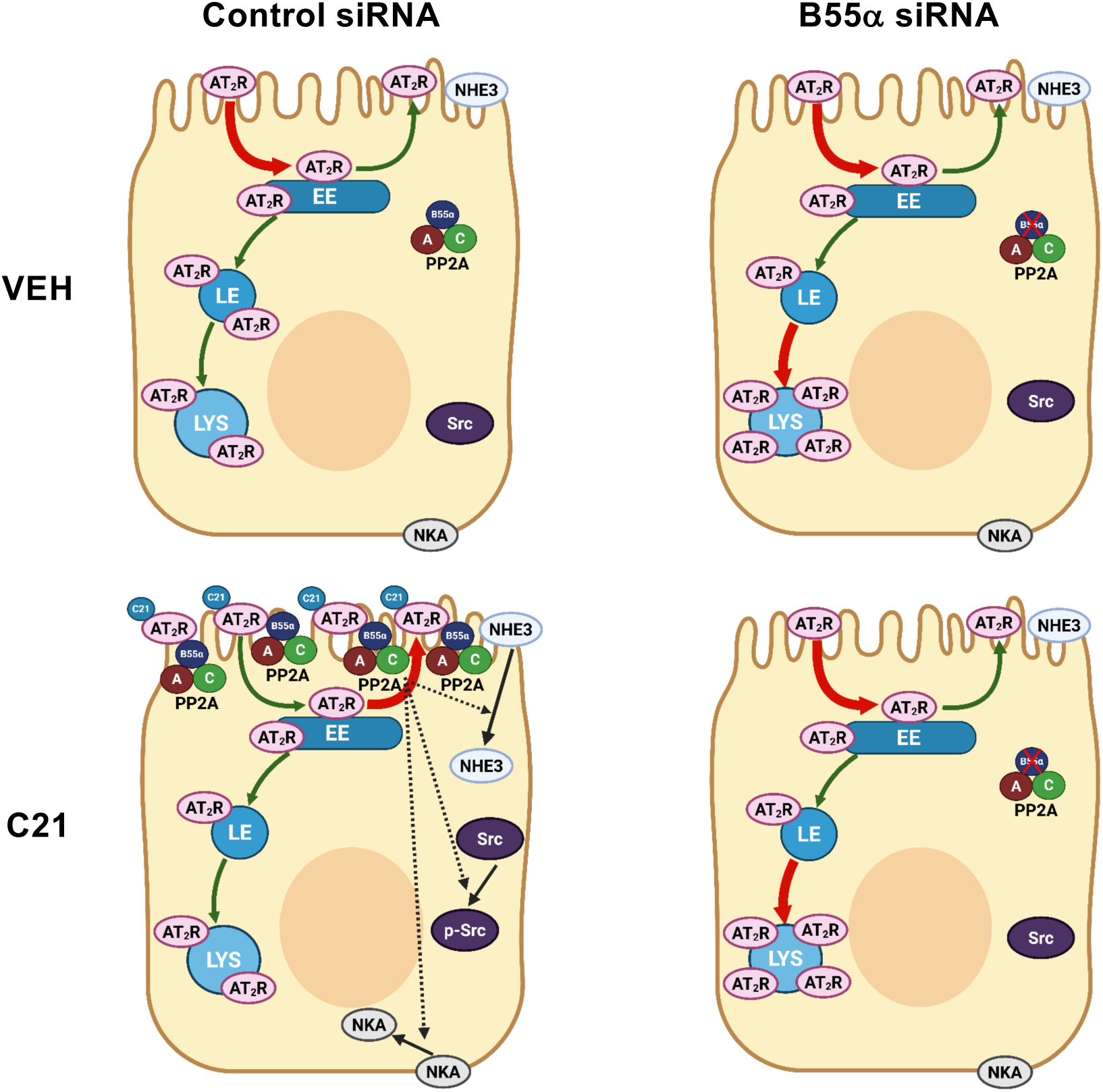
Schematic of AT_2_R signaling and trafficking without and with disruption after siRNA mediated knockdown of PP2A B55α. Under control siRNA/vehicle (VEH)-treated conditions, the AT_2_R recycles between the early endosomal (EE) compartment and the cell surface (apical brush border membranes) at a slow rate keeping AT_2_R at the cell surface low. A portion of the recycling AT_2_R in early endosomes traffics to late endosomes (LE) and lysosomes (LYS). AT_2_R agonist stimulation (C21) leads to increased AT_2_R binding to PP2A B55αC and AT_2_R/PP2A B55αC in apical brush border membranes, most likely due to a slow-down in AT_2_R internalization and/or promotion of AT_2_R recycling, allowing increased agonist binding and AT_2_R signaling to promote retrieval/internalization of the sodium transporters NHE3 and NKA and src phosphorylation and increase natriuresis. A smaller portion of the AT_2_R traffics to late endosomes during this phase. When B55α is knocked down, AT_2_R trafficking and signaling are disrupted and a large portion of the AT_2_R accumulates in lysosomes. The model was created using BioRender software.

In RPTCs of siRNA B55α-treated kidneys chronic AT_2_R internalization/recycling in the absence of B55α may promote AT_2_R trafficking to lysosomal compartments while decreasing it in early and late endosomal compartments. However, despite lysosomal targeting of the AT_2_R, it may not be degraded at an increased rate as total AT_2_R levels are similar between B55α-siRNA and control-siRNA treated RPTCs. Note, LAMP1 marks degradative as well as non-degradative lysosomes^41^. To which the AT_2_R localizes needs to be defined.

With our studies we cannot differentiate whether PP2A is only required for keeping the AT_2_R in the apical brush border membranes to allow further ligand binding and thereby promote signaling, or whether PP2A also plays additional roles in AT_2_R signaling. Previous studies identified PP2A as an intermediate player in AT_2_R signaling downstream of inhibitory G-proteins (Gi)^8^. Also, we previously showed that PP2A is necessary for cAMP and cGMP-mediated AT_2_R recruitment to the plasma membrane in immortalized cultured human RPTCs^13^ placing the point of action for PP2A downstream of the AT_2_R. The direct interaction of B55α with the AT_2_R in the apical brush border membranes as established in this study, suggest the PP2A AB55αC may act early in AT_2_R signaling. However, it is still possible that PP2A trimers containing B55α or other B-subunits play roles in further downstream signaling. Note, there was also an ∼2-fold increase in intracellular AT_2_R-PP2A B55α interaction in RPTCs (**Figure 1A and B**). Previous studies showed that AT_1_R internalization generates signaling platforms in endosomes^28^.

### siRNA-Mediated Knockdown of B55α in RPTCs

Studies on the physiological role(s) of B55α have been hindered by the finding that homozygous whole body B55α knockout mice are not viable^42^. Although siRNA-mediated knockdown of B55α has been performed in cells in culture as for example in HeLa cells^40^, our study using siRNA to knock down B55α *in vivo* to study a physiological B55α function is an exception. Our approach using local siRNA delivery in kidneys yielded B55α knockdown specifically in RPTCs allowing us to determine the role of B55α in AT_2_R-mediated natriuresis *in vivo*. We currently do not know how B55α was knocked down in renal proximal but not distal tubules. Possible reasons are application of siRNA into the renal cortex and effective uptake by RPTCs that normally serves the retrieval of glomerular filtrate components. We suggest that our approach can be applied to knocking down other proteins specifically in RPTCs.

In conclusion, PP2A B55α subunit directly binds AT_2_R and is required for normal AT_2_R trafficking and recruitment to RPTC apical brush border membranes for subsequent AT_2_R signaling to natriuresis. The molecular mechanisms of B55α action in AT_2_R signaling and trafficking need to be further investigated and substrate(s) for the AT_2_R-PP2A AB55αC complex determined. B subunit stabilization to increase PP2A activity has recently been targeted for cancer treatment^43^. Kidneys of pre-hypertensive SHR lack natriuretic and downstream signaling responses to intrarenal AT_2_R stimulation with Ang III or C21 concomitant with impaired AT_2_R recruitment to the apical brush border membranes and interaction with PP2A AB55αC^16^. And SHR have increased AT_2_R phosphorylation^37^. B55α in RPTCs may represent an attractive target for alleviating Na^+^ retention and thereby controlling blood pressure in hypertensive individuals.

## Nonstandard Abbreviations and Acronyms

Ang: angiotensin
AT_2_R: angiotensin type 2 receptor
BP: blood pressure
BSA: bovine serum albumin
C21: Compound 21
c-src: cellular tyrosine protein kinase src
DMEM: Dulbecco’s Modified Eagle Medium
EEA1: early endosome antigen 1
Giα: heterotrimeric Gi protein alpha subunit
GPCR: G protein-coupled receptor
HA: hemagglutinin
LAMP1: lysosomal associated membrane protein 1
MAP: mean arterial pressure
Na^+^: sodium
NHE3: sodium-hydrogen exchanger-3
NKA: sodium-potassium ATPase
PBS: phosphate buffered saline
PFA: paraformaldehyde
PLA: proximity ligation assay
PP2A: serine/threonine protein phosphatase 2A
RFU: relative fluorescence units
RI: renal interstitial
RPTC: renal proximal tubule cell
siRNA: small interfering RNA
SD: Sprague Dawley
TBS: tris-buffered saline
TBST^1^: tris-buffered saline with 0.1% Tween-20
TBST^2^: tris-buffered saline with 0.02% Tween-20
UNaV: urinary sodium excretion
VEH: Vehicle
WKY: Wistar Kyoto

## Acknowledgments

We thank Dr. Derek J. Taylor at Case Western Reserve University for generously providing purified PP2A B55α, and Dr. Tahir Hussain at the University of Houston for kindly supplying the AT_1_R antibody. We acknowledge the University of Virginia Research Histology Core for preparing the cryostat thin kidney sections for subsequent use in confocal fluorescence microscopy and Dr. Peng Xu at the University of Virginia for assisting with preparing the model in Figure 8.

## Novelty and Significance

### What is known

- Angiotensin type 2 receptor signaling induces natriuresis, counteracting sodium retention elicited by angiotensin type 1 receptor activation
- AT_2_R signaling promotes natriuresis in kidney proximal tubules by increasing its own cell surface expression and the retrieval/internalization of the sodium transporters NHE3 and NKA.
- Our previous research suggested protein phosphatase 2A (PP2A) functions downstream of the AT_2_R to promote natriuresis.

### What new information does this article contribute

- AT_2_R and the regulatory PP2A B55α subunit directly interact with each other, and the interaction increases dramatically in RPTC apical brush border membranes with AT_2_R stimulation.
- PP2A B55α is required for AT_2_R-elicited natriuresis, AT_2_R signaling and intracellular trafficking in RPTCs.
- This study introduces a unique approach to investigate a physiological B55α function, knocking down B55α *in vivo* specifically in renal proximal tubules cells using renal interstitial infusion of siRNA.

### Summary

Angiotensin type 2 receptor (AT_2_R) signaling promotes natriuresis in renal proximal tubule cells (RPTCs) counteracting sodium retention stimulated by AT_1_R. Early signaling events mediating AT_2_R-elicited natriuresis are unknown. Our previous research suggested protein phosphatase 2A (PP2A) functions downstream of the AT_2_R to promote natriuresis. In this study, we investigated the requirement for the regulatory PP2A subunit B55α in AT_2_R signaling and natriuresis. Our results demonstrate that the AT_2_R and PP2A B55α directly interact with each other, and the interaction increases dramatically in RPTC apical brush border membranes with AT_2_R stimulation. When B55α is knocked down *in vivo* in RPTCs, AT_2_R agonist-induced natriuresis is abolished concomitant with deficient AT_2_R redistribution to apical brush border membranes and AT_2_R signaling. Furthermore, AT_2_R subcellular localization shifts to a lysosomal compartment. Our results support a key role for PP2A B55α in regulating both AT_2_R subcellular trafficking and signaling to natriuresis in RPTCs. Kidneys of pre-hypertensive spontaneously hypertensive rats lack natriuretic and downstream signaling responses to intrarenal AT_2_R stimulation concomitant with impaired AT_2_R recruitment to apical brush border membranes and interaction with PP2A AB55αC. B55α in RPTCs may thus represent an attractive target for alleviating sodium retention and thereby controlling blood pressure in hypertensive individuals.

## SUPPLEMENTAL MATERIAL

### Supplemental Methods

#### *In vitro* Binding Assay with Purified HA-Tagged AT_2_R and B55α

HEK293 cells transfected with a hemagglutinin (HA)-tagged AT_2_R^18^ were cultured in Dulbecco’s Modified Eagle Medium with high glucose (Gibco, cat. no. 11995-040) containing 10% fetal bovine serum (Aldrich, cat. no. A2153) and 1% penicillin-streptomycin (Gibco, cat. no. 15140-122) at 37°C and 5% CO_2_. In initial experiments we observed that B55α endogenous to HEK293 cells co-immunoprecipitated with the HA-AT_2_R, and this interaction resisted washing with mild detergents. To purify the HA-AT_2_R from HEK293 cells without contamination with endogenous B55α, we prepared lysates using denaturing conditions as described^19^. Briefly, after washing cells on a 15 cm plate twice with cold PBS containing 0.5 mM MgCl_2_ and 0.9 mM CaCl_2_ (Gibco, cat. no. 14040-117), cells were lysed in 1.5 mL 4% SDS, 10 mM DTT, 100 mM Hepes, 300 mM NaCl, 1 mM EDTA pH 7.5 plus Halt protease inhibitors (Thermo Scientific, cat. no. 78430), and heated to 100°C for 5 min. Upon cooling, 150 µL 300 mM N-ethylmaleimide (25 mM final concentration) was added to alkylate cysteine residues. Following 5 min incubation, 12.5 mL 1.7% Thesit (non-ethylene glycol dodecyl ether, Sigma, cat. no. 88315) in 50 mM Hepes/150 mM NaCl pH 7.5 was added to solubilize proteins. The solution was centrifuged at 20,000 g for 30 min and the supernatant passed through a 0.45 μm filter (Gen Clone, cat. no. 25-246). HA antibodies conjugated to agarose (100 μL) (Sigma cat. no. A2095) were added to the filtrate. After a 1h incubation at room temperature while rocking on a shaking platform, beads were collected and washed 3 x with 1 mL 50 mM MOPS/125 mM NaCl pH 7.4. Pelleted beads with bound HA-AT_2_R were divided into 3 aliquots. One aliquot, serving as control for starting material, was incubated with 2X Laemmli sample buffer (BIO-RAD; cat. no.1610737) and the supernatant collected after centrifugation. The other two aliquots were incubated with purified PP2A B55α or binding buffer alone using a similar protocol as described^20^. Briefly, 100 μL binding buffer (20 mM Hepes, pH 7.5, 0.15 M NaCl, 0.1 mM MnCl_2_, 60 mM 2-mercaptoethanol, 0.1 mg/mL bovine serum albumin (BSA), 10% glycerol, 0.01% NP-40, 1 mM PMSF, 10 µL/mL Halt protease/phosphatase inhibitors (Thermo Scientific, cat. no. 78442) was added with or without 5 μg recombinant purified PP2A B55α (kindly provided by Dr. Derek J. Taylor at Case Western Reserve University). After a 90 min incubation at 4°C, beads were washed 3 x with 1 mL 20 mM Hepes, pH 7.5, 1.0 M NaCl, and 10% glycerol. After the final wash, proteins bound to the beads (HA-AT_2_R and associated proteins) were eluted in 2X Laemmli sample buffer. Samples were analyzed by immunoblotting as described^21^. Briefly, membranes after transfer and blocking were incubated overnight at 4°C with PP2A B subunit antibodies (Cell Signaling, cat. no. 2290; 1:1,000) in 5% BSA in TBST^1^. The membrane was subsequently incubated with HRP-conjugated secondary antibody (GE Healthcare, cat. no. NA934, 1:2,500) in 5% milk TBST^1^ for 2h at room temperature. Signals were detected using chemiluminescence (ECL Select Western Blotting Detection Reagent, Cytiva, cat. no. 2235) and the BIO-RAD ChemiDoc MP Imaging System. Signal intensities were analyzed with ImageJ software. The membrane was stripped for 15 min using Restore Western blot stripping buffer (Thermo Scientific; cat. no. 21059), blocked in 5% milk TBST^1^ for 1h at room temperature, and then incubated with AT_2_R antibody (Abcam; Cat # ab92445; 1:1,000) in 5% BSA TBST^1^ overnight at 4°C, followed by secondary antibody, and detection as described above.

## Supplemental Figures

**Figure S1.**
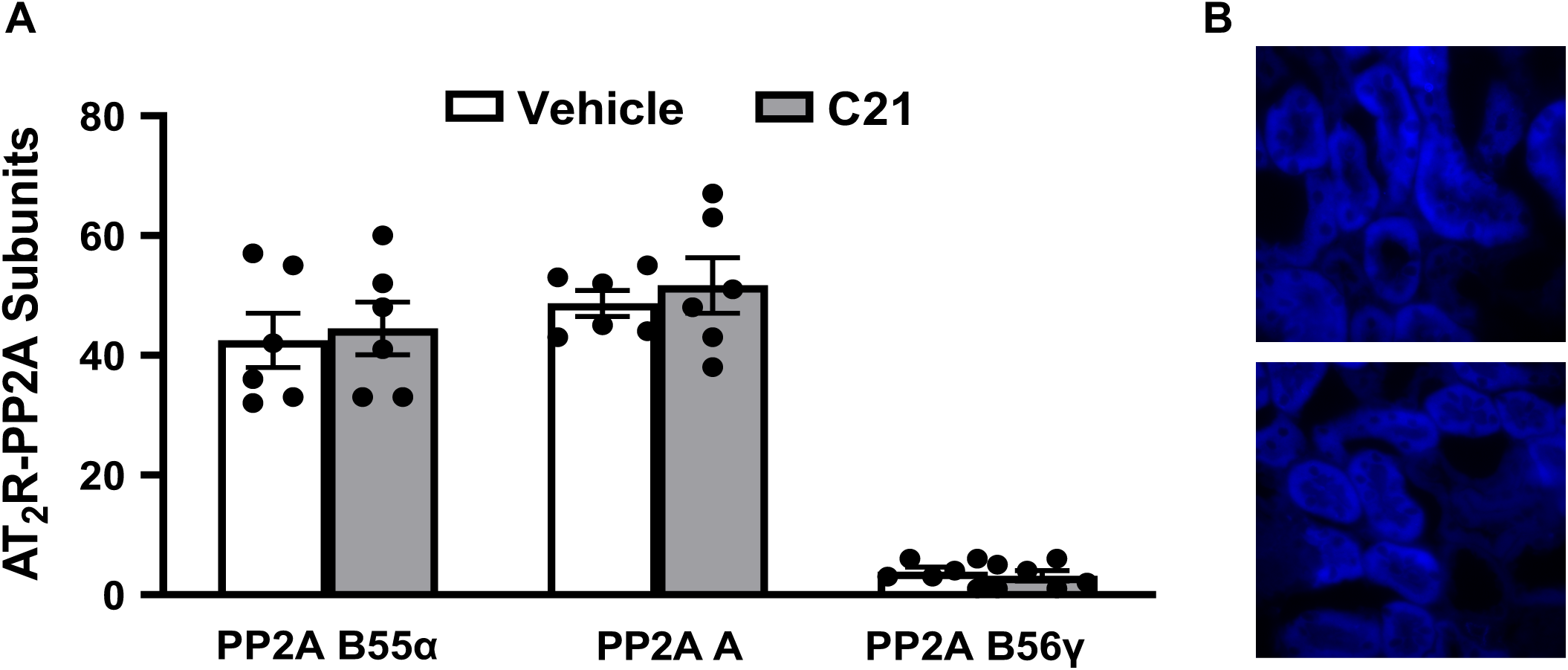
Kidney sections were isolated from Wistar Kyoto rats after renal interstitial (RI) infusion of vehicle or compound 21 (C21). **A.** Interactions between the AT_2_R and PP2A subunits B55α, A, and B56γ outside renal proximal tubules as identified by proximity ligation assay. Numbers represent the number of discrete green dots outside proximal tubules. Data are shown as mean±SE (n=6) and were analyzed using one-way ANOVA analysis. **B.** Negative controls for proximity ligation assays did not yield any green fluorescent dots. Kidney sections were subjected to PLA incubation steps using non-specific mouse IgG (Santa Cruz; cat. no. sc-2025) together with AT_2_R rabbit antibodies (Santa Cruz H-143; cat. no. sc-9040) (top image) or non-specific rabbit IgG (Sigma; cat. no. I5006) and B55α mouse antibodies (Santa Cruz; cat. no. sc-81606) (bottom image). Autofluorescence (blue, DAPI channel) identifies RPTCs.

**Figure S2.**
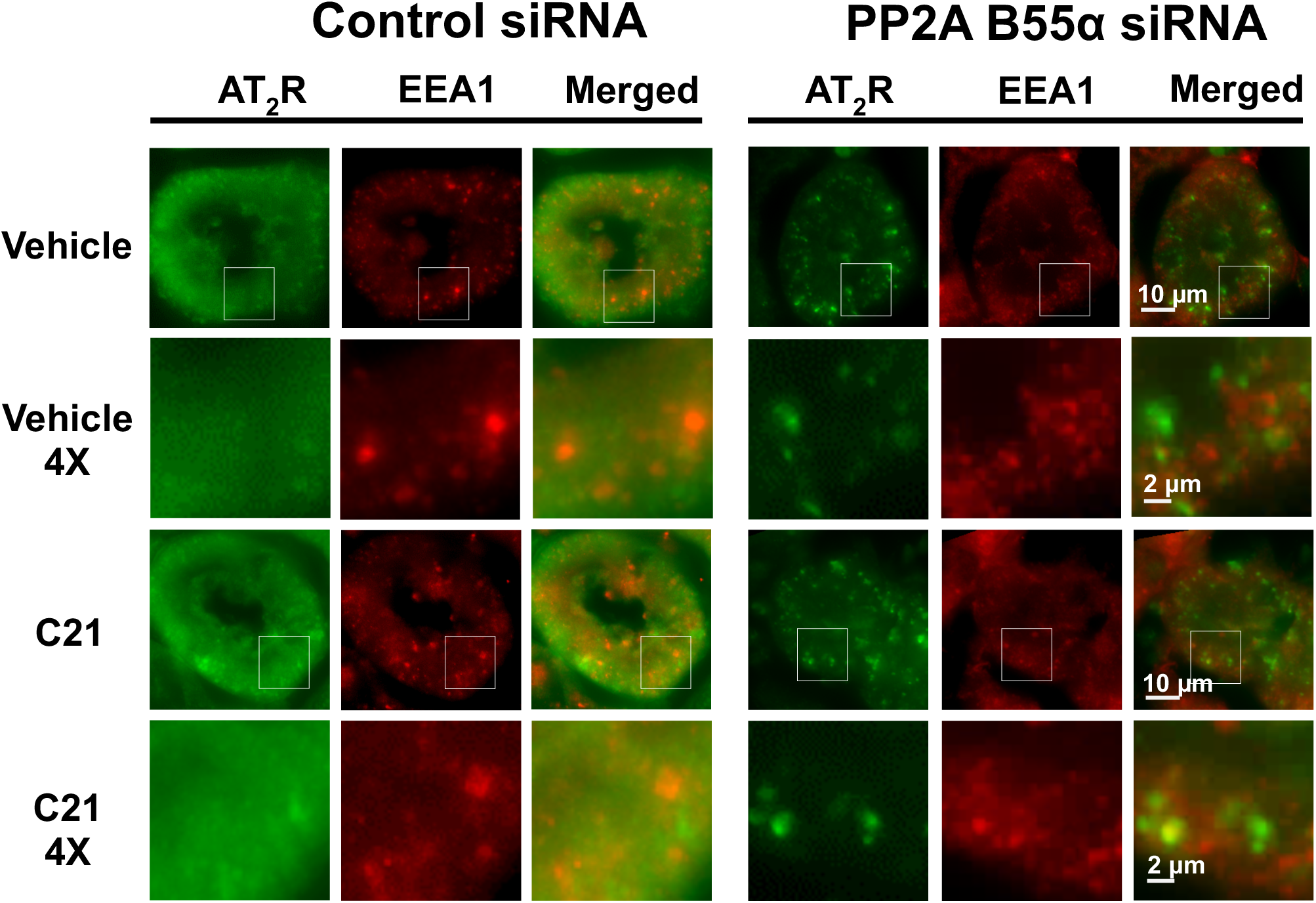
Co-localization of the AT_2_R with EEA1 in renal proximal tubule cells. Sprague-Dawley rats were treated with control siRNA and PP2A B55α siRNA by renal interstitial (RI) infusion 48 hours prior to RI infusion of vehicle or compound 21 (C21) and kidney sections analyzed by confocal microscopy analysis. Kidney sections were co-stained with AT_2_R and EEA1 antibodies and representative images of renal proximal tubule cells are shown for each condition and each antibody separately and together (merged) as labeled. Squares outlined in images at 1X were magnified (4X) to better show co-localization of AT_2_R with EEA1.

**Figure S3.**
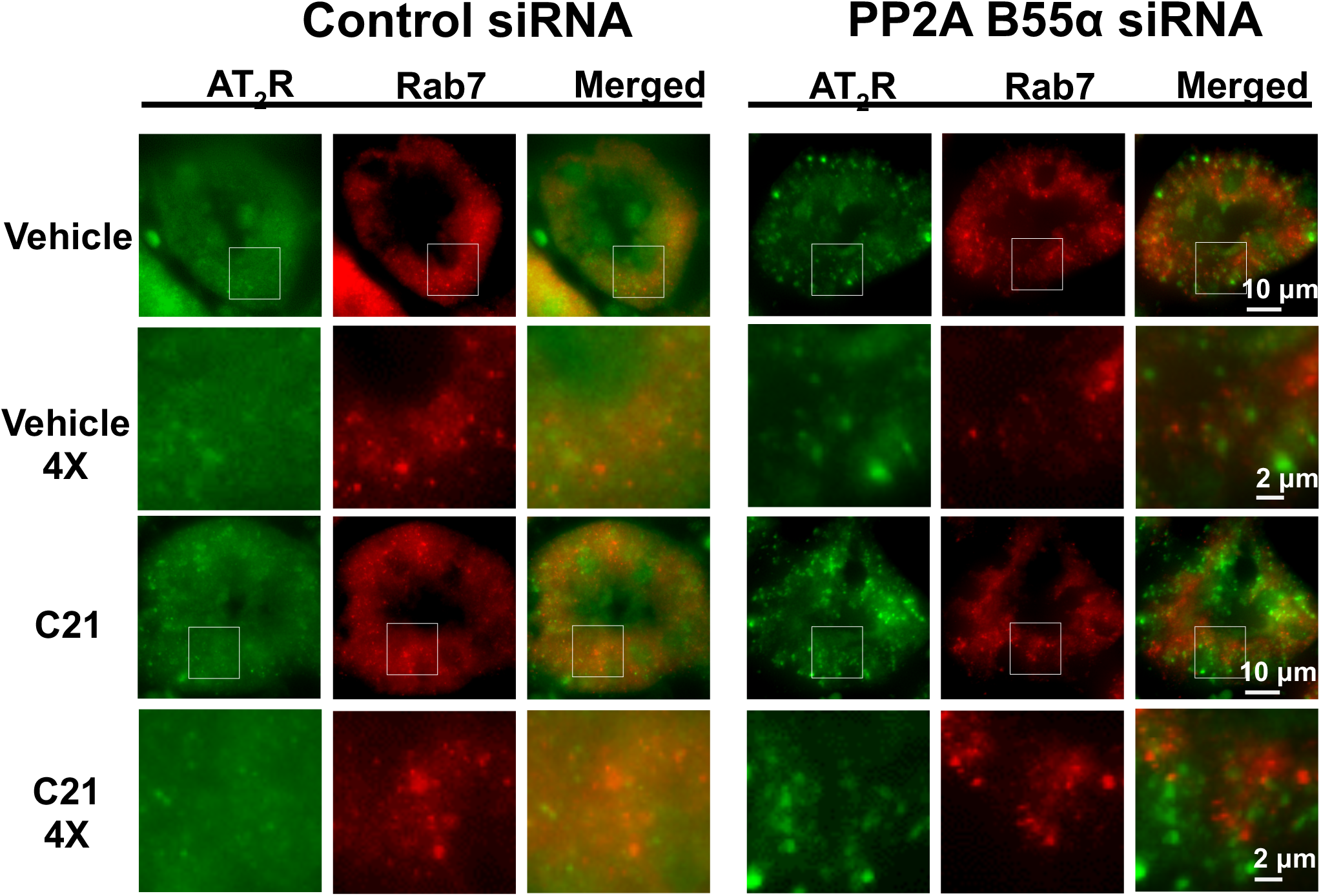
Co-localization of the AT_2_R with Rab7 in renal proximal tubule cells. Sprague-Dawley rats were treated with control siRNA and PP2A B55α siRNA by renal interstitial (RI) infusion 48 hours prior to RI infusion of vehicle or compound 21 (C21) and kidney sections analyzed by confocal microscopy analysis. Kidney sections were co-stained with AT_2_R and Rab7 antibodies and representative images of renal proximal tubule cells are shown for each condition and each antibody separately and together (merged) as labeled. Squares outlined in images at 1X were magnified (4X) to better show co-localization of AT_2_R with Rab7.

**Figure S4:**
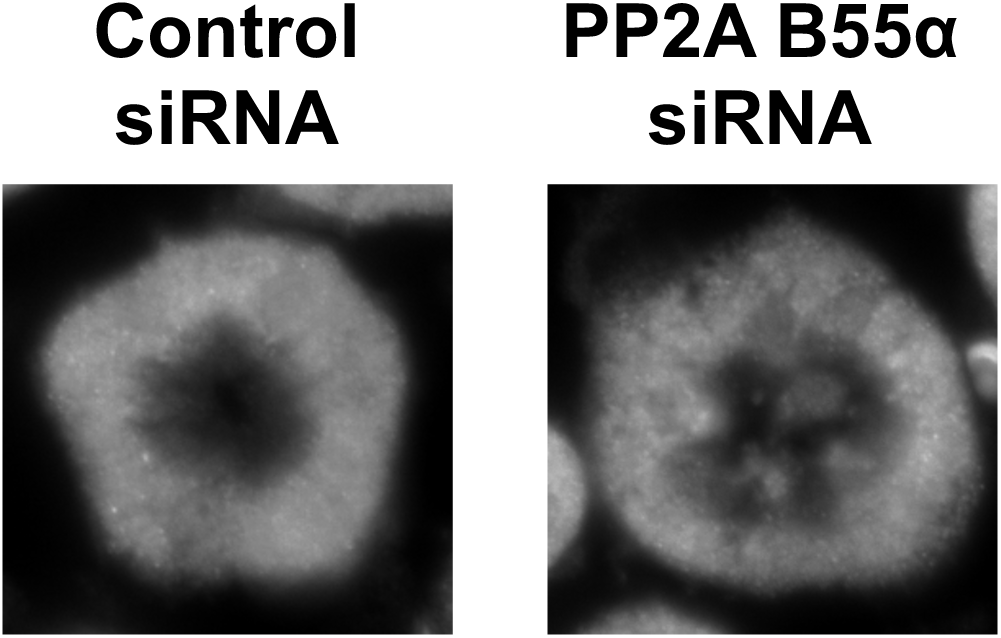
AT_1_R localization in renal proximal tubule cells. Sprague-Dawley rats were treated with control siRNA and PP2A B55α siRNA by renal interstitial (RI) infusion 48 hours prior to RI infusion of vehicle and kidney sections analyzed by confocal microscopy analysis. Kidney sections were stained with AT_1_R antibodies and representative images of renal proximal tubule cells are shown for control siRNA and PP2A B55α siRNA.

**Related to Figure 2:**
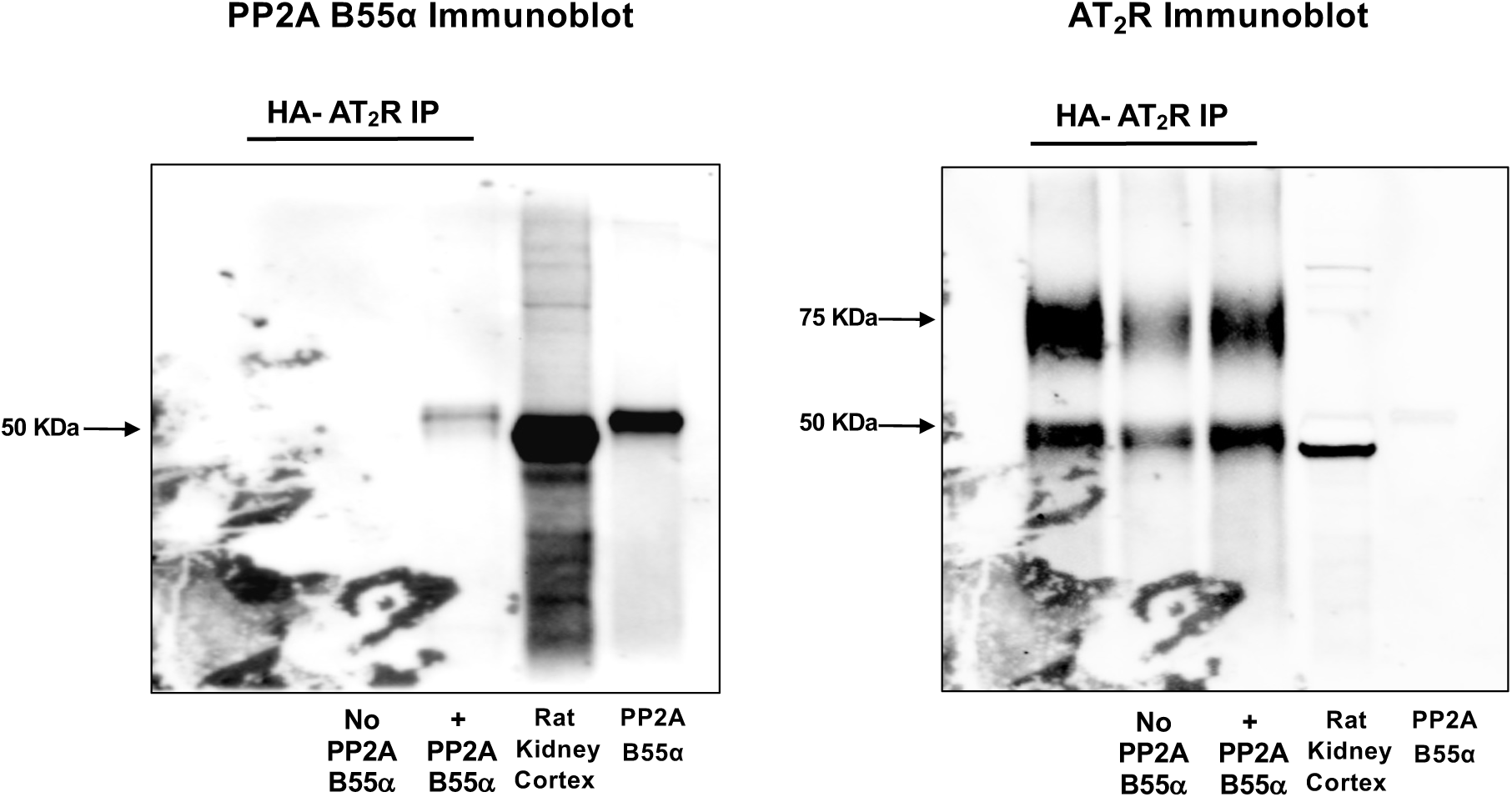
Uncut Immunoblots

**Related to Figures 3, 5, 6, S2 and S4:**
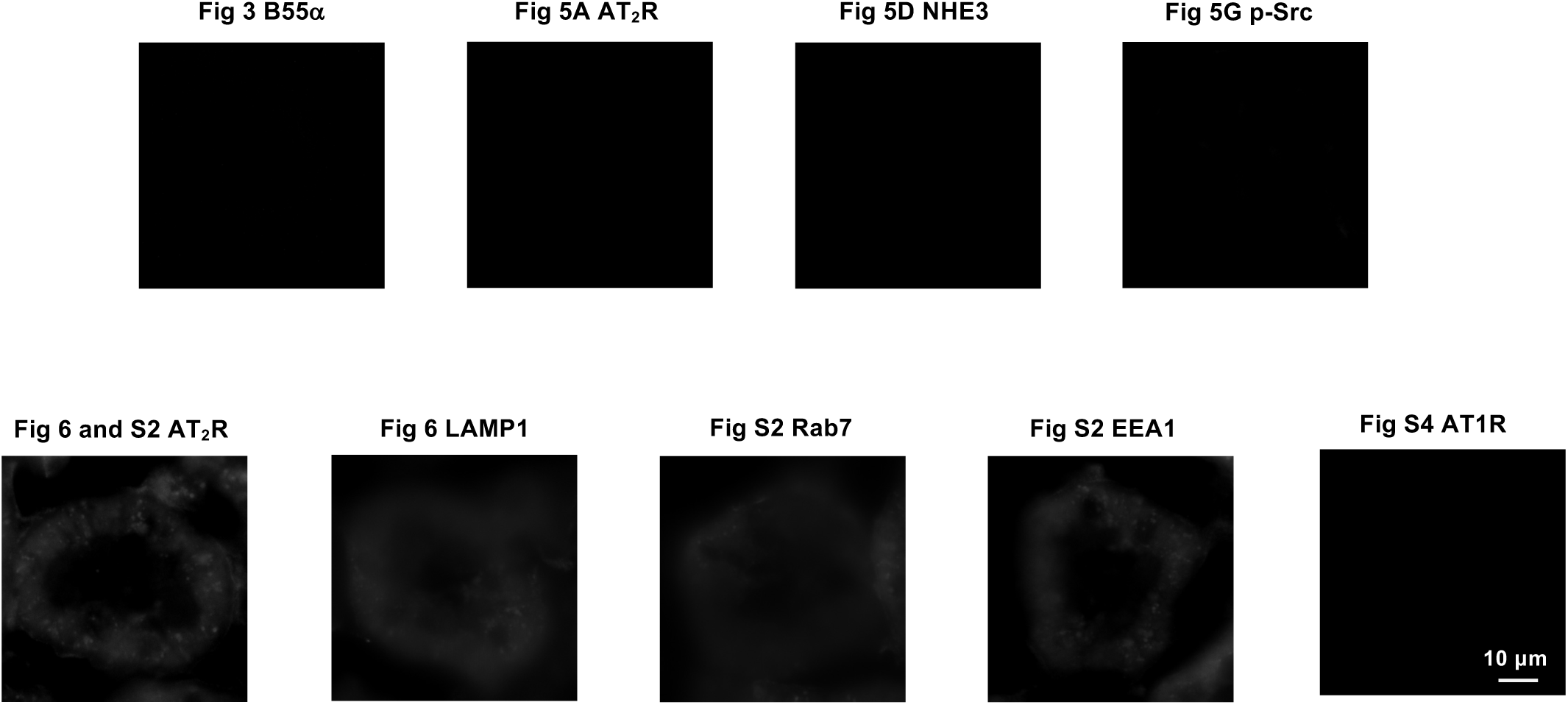
Confocal Background Staining (no respective primary antibodies)

## Major Resources Table

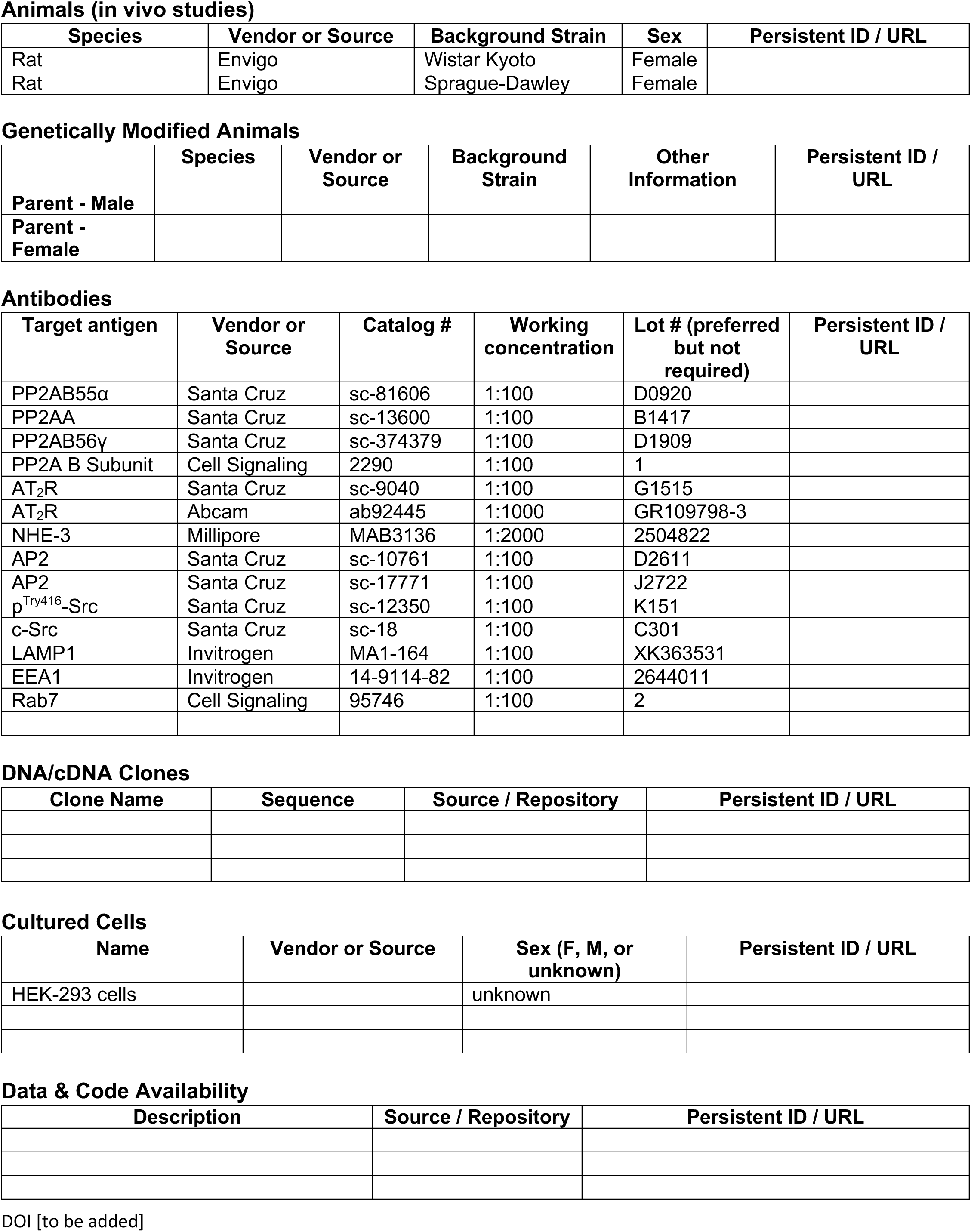

Animals (in vivo studies)
| Species | Vendor or Source | Background Strain | Sex | Persistent ID / URL |
| --- | --- | --- | --- | --- |
| Rat | Envigo | Wistar Kyoto | Female |  |
| Rat | Envigo | Sprague-Dawley | Female |  |

Genetically Modified Animals
|  | Species | Vendor or Source | Background Strain | Other Information | Persistent ID / URL |
| --- | --- | --- | --- | --- | --- |
| Parent - Male |  |  |  |  |  |
| Parent - Female |  |  |  |  |  |

Antibodies
| Target antigen | Vendor or Source | Catalog # | Working concentration | Lot # (preferred but not required) | Persistent ID / URL |
| --- | --- | --- | --- | --- | --- |
| PP2AB55α | Santa Cruz | sc-81606 | 1:100 | D0920 |  |
| PP2AA | Santa Cruz | sc-13600 | 1:100 | B1417 |  |
| PP2AB56γ | Santa Cruz | sc-374379 | 1:100 | D1909 |  |
| PP2A B Subunit | Cell Signaling | 2290 | 1:100 | 1 |  |
| AT<sub>2</sub>R | Santa Cruz | sc-9040 | 1:100 | G1515 |  |
| AT<sub>2</sub>R | Abcam | ab92445 | 1:1000 | GR109798-3 |  |
| NHE-3 | Millipore | MAB3136 | 1:2000 | 2504822 |  |
| AP2 | Santa Cruz | sc-10761 | 1:100 | D2611 |  |
| AP2 | Santa Cruz | sc-17771 | 1:100 | J2722 |  |
| p<sup>Try416</sup>-Src | Santa Cruz | sc-12350 | 1:100 | K151 |  |
| c-Src | Santa Cruz | sc-18 | 1:100 | C301 |  |
| LAMP1 | Invitrogen | MA1-164 | 1:100 | XK363531 |  |
| EEA1 | Invitrogen | 14-9114-82 | 1:100 | 2644011 |  |
| Rab7 | Cell Signaling | 95746 | 1:100 | 2 |  |

DNA/cDNA Clones
| Clone Name | Sequence | Source / Repository | Persistent ID / URL |
| --- | --- | --- | --- |

Cultured Cells
| Name | Vendor or Source | Sex (F, M, or unknown) | Persistent ID / URL |
| --- | --- | --- | --- |
| HEK-293 cells |  | unknown |  |

Data & Code Availability
| Description | Source / Repository | Persistent ID / URL |
| --- | --- | --- |

DOI [to be added]

